# Intrinsically disordered regions regulate RhlE RNA helicase functions in bacteria

**DOI:** 10.1101/2024.01.12.575364

**Authors:** Stéphane Hausmann, Johan Geiser, George Edward Allen, Sandra Amandine Marie Geslain, Martina Valentini

## Abstract

RNA helicases—central enzymes in RNA metabolism— often feature intrinsically disordered regions (IDRs) that enable phase separation and complex molecular interactions. In the bacterial pathogen *Pseudomonas aeruginosa*, the non-redundant RhlE1 and RhlE2 RNA helicases share a conserved REC catalytic core but differ in C-terminal IDRs. Here, we show how the IDR diversity defines RhlE RNA helicase specificity of function. Both IDRs facilitate RNA binding and phase separation, localizing proteins in cytoplasmic clusters. However, RhlE2 IDR is more efficient in enhancing REC core RNA unwinding, exhibits a greater tendency for phase separation, and interacts with the RNase E endonuclease, a crucial player in mRNA degradation. Swapping IDRs results in chimeric proteins that are biochemically active but functionally distinct as compared to their native counterparts. The REC_RhlE1_-IDR_RhlE2_ chimera improves cold growth of a *rhlE1* mutant, gains interaction with RNase E and affects a subset of both RhlE1 and RhlE2 RNA targets. The REC_RhlE2_-IDR_RhlE1_ chimera instead hampers bacterial growth at low temperatures in the absence of RhlE1, with its detrimental effect linked to aberrant RNA droplets. By showing that IDRs modulate both protein core activities and subcellular localization, our study defines the impact of IDR diversity on the functional differentiation of RNA helicases.

## Introduction

RNA molecules, regardless of their length, have the capacity to adopt multiple conformations. However, many of these conformations, although thermodynamically stable under physiological conditions, may not be functional [1]. To ensure that RNA molecules attain their correct functional conformations and to resolve non-functional structures, RNA helicases play a crucial role in nearly all species across various domains [2]. Often referred to as the ‘molecular motors’ of the cell, RNA helicases harness the energy derived from ATP binding and hydrolysis to unwind RNA duplexes, reshape RNA secondary structures, and modulate ribonucleoprotein complexes [3, 4].

The largest family of RNA helicases is the DEAD-box proteins [5]. They are characterized by a globular catalytic core (called REC core) composed of two RecA-like domains, homologous to the bacterial RecA protein, connected by a short flexible linker [6]. Within the REC core, there are 12 highly conserved sequence motifs (including the D-E-A-D motif giving the name to the family) that are critical for ATP or RNA binding, inter-subunit interactions, and the ATPase activity of the protein [7, 8]. RNA unwinding is the action of DEAD-box RNA helicases that has been most extensively characterized at the molecular level [3]. Typically, ATP binds to a conserved pocket at the interface between the two RecA domains, leading to a conformational change that exposes a positively charged region capable of interacting with the RNA phosphate backbone. Binding to RNA induces a local distortion in the RNA structure, rendering it incompatible with an RNA duplex. This leads to the retention of one RNA single stand within the core, while the second strand dissociates. Subsequent ATP hydrolysis and the release of ADP allows the release of the protein-bound RNA strand from the enzyme, which recovers its original conformation and is regenerated for another round of RNA unwinding [reviewed in[2, 3, 5, 7-10].

In the same organism, multiple RNA helicases can be present, each exhibiting distinct and specific functions [2]. Since the REC core interacts with RNA in a sequence-independent manner, it has been shown that additional domains, known as accessory domains, can confer specificity to RNA helicases [9-11]. Many DEAD-box RNA helicases feature intrinsically disordered regions (IDRs) at their N- and/or C-terminal ends, either in lieu of or alongside structured accessory domains [12]. IDRs are protein regions characterized by the absence of a stable three-dimensional structure while being composed of flexible conformational ensembles. Their inability to adopt a well-defined conformation is primarily attributed to their high content of polar and charged residues (specifically, G, R, Q, S, E, and K) and the insufficient presence of hydrophobic amino acids required for cooperative folding, as compared to structured domains [13]. Once thought of as mere links between structured domains, IDRs play instead active roles in protein function, including enabling interaction with multiple partners or driving liquid-liquid phase separation (LLPS)[14-18]. LLPS is an equilibrium process through which biomolecules (proteins or RNAs) segregate in a solution by weakly associating with each other and form condensed micrometer-sized liquid droplets [19]. Within the cell, LLPS can induce the formation of subcellular microcompartments, called biomolecular condensates, which can favour or restrict specific biochemical reactions [20]. Several RNA helicases that possess IDRs have been shown to undergo LLPS [21, 22]. Nonetheless, the rapid evolution and diversity of IDRs pose a challenge in understanding their contribution to the function of RNA helicases and make their role difficult to predict [23, 24].

Our research has focused on DEAD-box RNA helicases in the opportunistic pathogen *Pseudomonas aeruginosa*, particularly on two homologous proteins named RhlE1 and RhlE2 [25, 26]. The RhlE RNA helicase group is prevalent in Proteobacteria, often with multiple copies within a single genome possibly originated from gene duplication in a Proteobacterial ancestor [27, 28]. We have previously shown that in *P. aeruginosa rhlE* gene duplication did not lead to redundancy but rather sub-functionalization. Both RhlE1 and RhlE2 play crucial roles in bacterial growth at low temperatures, but their lack of cross-complementation suggested that their functions were neither overlapping nor redundant. While RhlE1 primarily supports cold growth, RhlE2 has broader impacts on various phenotypes of *P. aeruginosa*, including virulence. Furthermore, RhlE2 interacts with the RNase E endonuclease, a major component of the RNA degradation machinery in many bacteria, and influences RNA stability [25, 29]. The interaction occurred via the RhlE2 C-terminal extension (CTE), and it is mediated or stabilized by RNA [25].

Given that many of the differences between the sequences of RhlE1 and RhlE2 are concentrated in their CTEs, we sought to characterize how the CTEs contribute to the specific functions of the proteins. Sequence analysis predicted that both CTEs were intrinsically disordered although with varying length and amino acid composition (Figure S1). In this study, we conducted a systematic examination of the REC-CTE modules of RhlE1 and RhlE2 individually, in native combinations or by constructing chimeras. The analysis included the characterization of phenotypes and transcriptomes, assessment of protein-protein interactions, evaluation of ATPase activity, RNA binding, RNA unwinding capacity, properties related to liquid-liquid phase separation (LLPS), and the formation of biomolecular condensates. Collectively, the data reveal that the distinct functional roles of RhlE1 and RhlE2 are driven by the differential modulation of REC core activities by their respective IDRs, demonstrating a co-dependent relationship between these modules.

## Results

### Role of IDRs of RhlE RNA helicases in sustaining *P. aeruginosa* growth at cold temperatures

We have previously observed that RhlE RNA helicases have clade-specific, rapidly evolving C-terminal extensions (CTEs). In *P. aeruginosa*, RhlE1 and RhlE2 share a conserved REC core but possess different CTEs in terms of length and amino acid composition. RhlE1 has a 70-residue poly-lysine stretch, while RhlE2 has basic 255-residue CTE with a poly-glutamine stretch (Figure S1A). Both RhlE1 and RhlE2 CTEs, along with those of other RhlE RNA helicases, are predicted to be highly disordered (Figure 1A and Figure S1B). To understand the role of the CTEs in the function of RhlE proteins, we constructed strains expressing either the REC core or CTE separately (named R1 or C1 for REC and CTE of RhlE1 and R2 and C2 for RhlE2, respectively) and two chimeras in which the CTEs of RhlE1 and RhlE2 were swapped (named R1C2 and R2C1, Figure 1B). The protein variants were expressed under the control of an arabinose inducible promoter to achieve comparable expression levels in all strains (Figure S2). We previously observed that both *rhlE1* and *rhlE2* mutants were affected in growth at 16°C as compared to the wild-type strain [25]. Cold-sensitivity phenotype is a most common phenotype of bacterial RNA helicase mutants, likely due to their role in counteracting the increased stability of mRNA duplexes at low temperatures, thereby promoting mRNA translatability and turnover [30, 31]. Therefore, we evaluated the *in vivo* functionality of each REC/CTE protein module and RhlE variants by their ability to restore growth of the Δ*rhlE1* (E1D), Δ*rhlE2* (E2D) or Δ*rhlE1*Δ*rhlE2* (E1DE2D) mutant at 16°C as compared to the native proteins (Figure 1C and Figure S3). Expression of the R1 (i.e., RhlE1^1-392^) or R2 domain (i.e., RhlE2^1-384^) and C1 or C2 alone (i.e., RhlE1^379-449^ and RhlE2^384-634^ respectively) had no impact on growth in any genetic background (Figure S3A and B), suggesting that no single RNA helicase module is functional on its own. The expression on the R1C2 chimera slightly but significantly improved the growth of E1D and E1DE2D mutants (Figure 1C, compare column 4 vs 2 and 13 vs 10), suggesting some retention of RhlE1 activity. A similar effect was seen when testing planktonic growth at 16 °C (Figure S3C). The effect of R1C2 in a E2D background was not evident, but RhlE2 seems to affect colony size rather than CFU number (Figure 1C, compare column 8 vs 6). The R2C1 chimera was unexpectedly deleterious for growth of the E1D and E1DE2D mutants (Figure 1C column 5 and 14, and Figure S3C) but did not negatively impact growth in an E2D background, suggesting an interference with RhlE1 regulation in the absence of its native form. Since the R2C1 chimera impaired growth only at 16°C and not at 37°C (Figure S3D), we conclude that the inhibitory effect is not unspecific, but related to the regulation of RNA metabolism at low temperatures.

**Figure 1.**
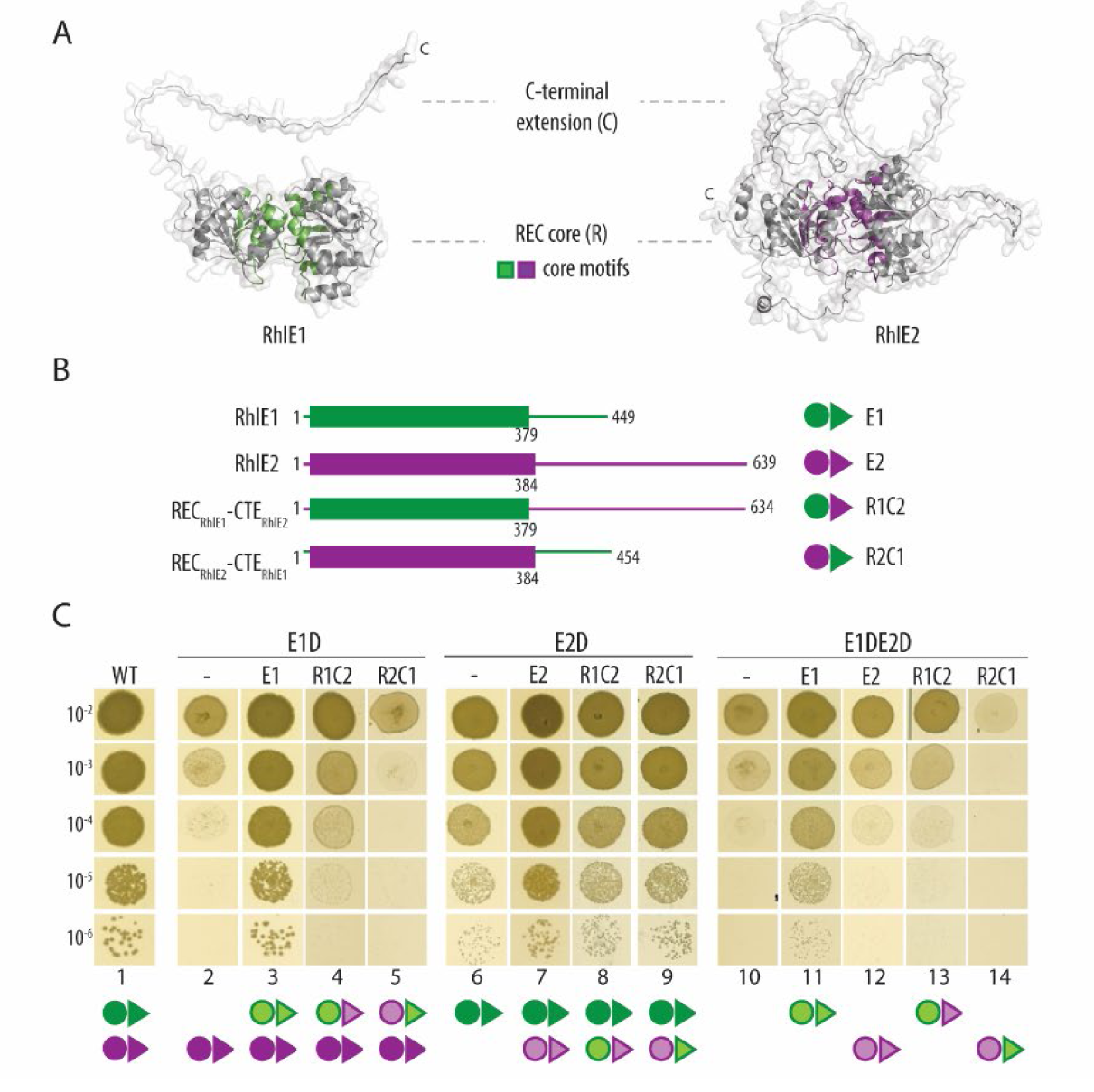
Interchangeability of REC and CTE modules RhlE1 and RhlE2 for cold adaptation. **(A)** Alphafold predicted structure of *P. aeruginosa* RhlE1 and RhlE2 [71, 72]. Core motifs within the catalytic REC core of RhlE1 and RhlE2 are coloured in green and violet, respectively. **(B)** Linear representation of RhlE1 (E1), RhlE2 (E2), REC_RhlE1_-CTE_RhlE2_ chimera (R1C2) and REC_RhlE2_-CTE_RhlE1_ (R2C1) chimera. The REC catalytic domain is represented as coloured box and the C-terminal extension (CTE) as line. **(C)** Growth on cold (16 °C) of wild type PAO1 (WT), Δ*rhlE1* mutant (E1D), Δ*rhlE2* mutant (E2D) and Δ*rhlE1*Δ*rhlE2* mutant (E1DE2D), along with the corresponding strains expressing either 3xFLAG-tagged E1, E2, R1C2, or R2C1 under the control of an AraC-P_BAD_ promoter using mini-Tn7 constructs (see Table S1). Strains were grown on agar LA plates with 0.2% arabinose as indicated. Experiments were performed in biological independent triplicates as described in Materials and Methods. Figure in panel C illustrates a representative LA plate.

Collectively, these results demonstrate that IDR swapping alters the functionality of RNA helicases. To further understand the activities of the chimeras and the roles of the CTEs/IDRs, we performed (i) a biochemical characterization of the chimeras and separate domains relative to the native proteins, (ii) an assessment of their liquid-liquid phase separation (LLPS) behaviour, and (iii) a transcriptome analysis of strains expressing R1C2 and R2C1 in the E1D, E2D, or E1DE2D backgrounds, as outlined in subsequent sections.

### Differential abilities of the RhlE1 and RhlE2 CTEs/IDRs in promoting RNA unwinding

We have previously demonstrated that both RhlE1 and RhlE2 possess an RNA-dependent ATPase activity and an ATP-dependent RNA unwinding activity [25]. To evaluate the influence of the IDRs on the protein activities, we performed quantitative assessments of ATPase activity, RNA binding, and RNA unwinding for truncated proteins as well as for the R1C2 and R2C1 chimeras, compared to the native full-length RhlE1 (E1) and RhlE2 (E2) proteins. We designed two truncations for E1 (E1^1-379^ and E1^1-392^) and one for E2 (E2^1-384^). The design of the shortest truncations (E1^1-379^ and E2^1-384^) was based on the construction of the chimeras (Figure 1B). All variants were successfully expressed in *E. coli* as N-terminal His10-Smt3-tagged fusion and purified by adsorption to nickel-agarose (Figure S4). However, the E1^1-379^ truncated protein exhibited lower solubility and increased susceptibility to freeze-thaw cycles, leading us to design and purify the slightly longer variant E1^1-392^, which still lacks the disordered lysine-rich basic region present in the CTER of RhlE1, but shows improved stability. Results for both truncations are presented for the sake of completeness (for E1^1-379^ data see Figure S5).

In the presence of 5 µM of a 5’ overhang RNA duplex (5’ovh_dsRNA_26/14_), the RNA-dependent ATPase activity of both E1^1-392^ closely resembled those of the E1, respectively (17.6 vs 18.2 min^-1^) while E1^1-379^ activity is slightly lower (8.6 min^-1^) yet still on a similar range of E1 (Figure 2 A and B and S5A). The same was observed for E2^1-384^ and E2 full-length ATPase activities (78.5 vs 60.9 min^-1^, Figure 2 C and D). These observations indicate that the ATPase activity of the REC cores is not regulated by the CTEs. In line with this finding, the R1C2 chimera exhibited similar activity (27.4 min^-1^) to that of E1, and R2C1 (57.4 min^-1^) mirrored the activity of E2 (Figure 2 E and F).

**Figure 2:**
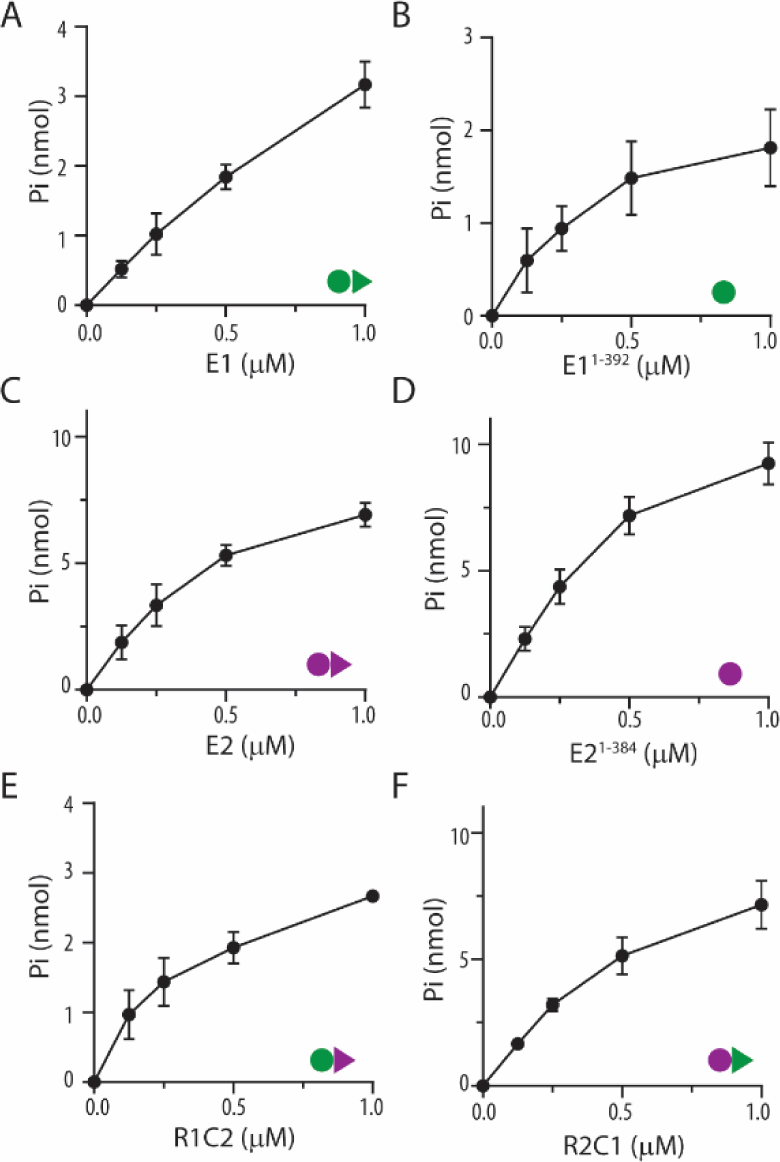
RNA-dependent ATPase activity of different RhlE proteins and their variants. Reaction mixtures (15 µl) containing 50 mM HEPES (pH 7.2-7.5), 1 mM DTT, 50 mM KGlu, 2 mM MgGlu, 1 mM [γ -32P] ATP, 5 µM duplex RNA (5’ovh_dsRNA26/14) and either **(A)** RhlE1 (E1), **(B)** RhlE1^1-392^ (E1^1-392^), **(C)** RhlE2 (E2), **(D)** RhlE2^1-384^ (E2^1-384^), **(E)** R1C2 or **(F)** R2C1 chimera were incubated for 15 min at 37 °C. Pi release was determined as described in Materials and Methods and was plotted as a function of protein concentration. Data are the average of three independent experiments with error bars representing standard deviation. Specific activity for each protein was calculated from the slope of the titration curve in the linear range and values are reported in Table 1.

**Table 1:** Biochemical characterization of RhlE proteins.

| Protein | ATPase activity<br>(min<sup>-1</sup>)* | RNA affinity<br>(nM)** | Unwinding<br>activity (min<sup>-1</sup>)*** |
| --- | --- | --- | --- |
| RhlE1 | 18.2 | 1.7 | 0.0317 |
| RhlE1<sup>1-392</sup> (R1) | 17.6 | 73 | 0.0003 |
| RhlE2 | 60.9 | <0.8 | 0.4368 |
| RhlE2<sup>1-384</sup> (R2) | 78.5 | 7.5 | 0.0105 |
| R1C2 | 27.4 | 1.1 | 0.1041 |
| R2C1 | 57.4 | 1.5 | 0.0209 |

\*: see Figure 2, \*\*: see Figure 3, \*\*\*: see Figure 4.

To assess the ability of the proteins to bind RNA, we used the fluorescently (5-carboxyfluorescein [5-FAM]) labelled 5’ovh_dsRNA_26/14_ and we performed fluorescence polarization assays in the presence of the ATP analogue adenosin-5′-(β,γ-imido)triphosphate (AMPPNP) and Mg^2+^ ions (Figure 3). The data were fitted as described in the Methods sections, and dissociation constants (Kd) were calculated based on the assumption that a monomeric protein binds to one substrate in a simple bimolecular binding reaction. Full-length E2 exhibited a slightly higher affinity for RNA compared to E1, with Kd values of <0.8 nM and 1.7, respectively (Figure 3 A and C). When the CTE was removed, the affinity of E1^1-392^ for RNA decreased by 43-fold (Kd = 73 nM), and E1^1-379^ exhibited a 57-fold decrease in affinity (Kd = 97 nM) compared to E1 full-length (Figure 3 A and B and S5B). In the case of E2^1-384^, its affinity for RNA decreased by at least 9-fold (Kd = 7.5 nM) when compared to the full-length E2 (Figure 3 C and D). On the other hand, the chimeras displayed similar affinities to RNA as the native proteins, with R1C2 having a Kd of 1.1 nM and R2C1 a Kd of 1.5 nM (Figure 3 E-F). Altogether, these findings suggest the CTEs promote RNA binding and reveal the interchangeable nature of the IDRs in enhancing RNA binding. To confirm this, we tested RNA binding affinities of E1 CTE (C1) and E2 CTE (C2). Due to technical limitations, we couldn’t determine the dissociation constant (Kd) of C1 and C2 with 5’ovh_dsRNA26/14, likely due to minimal size differences between the labelled RNA and the CTEs. To address this, we used a slightly smaller RNA, a 31-mer molecule with a hairpin (refer to Materials and methods). With this approach, we found that C1 and C2 exhibited similar Kd values compared to their full-length counterparts (Figure S7).

**Figure 3.**
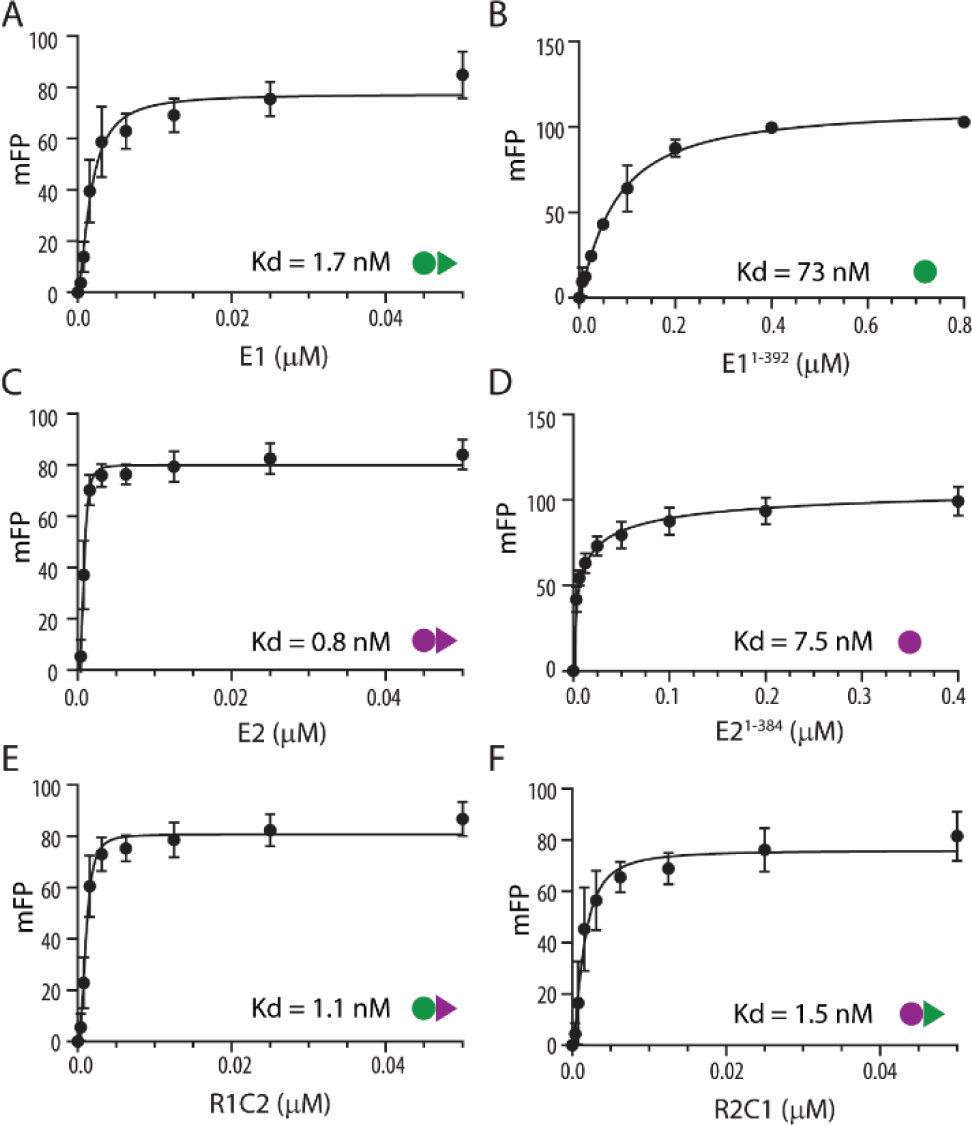
RNA binding of different RhlE proteins and their variants measured by fluorescence polarization assays. Reaction mixtures (20 µl) containing 50 mM HEPES (pH 7.2-7.5), 1 mM DTT, 50 mM KGlu, 2 mM MgGlu, 1 mM non-hydrolysable ATP analogue, 0.5 nM 5’-FAM-labeled duplex RNA (5’ovh_dsRNA26/14) and either **(A)** RhlE1 (E1), **(B)** RhlE1^1-392^ (E1^1-392^), **(C)** RhlE2 (E2), **(D)** RhlE2^1-384^ (E2^1-384^), **(E)** R1C2, **(F)** R2C1 protein as specified were incubated for 15 min at 25 °C. mFP was determined as described in Material and Methods and was plotted as of function of protein concentration. Data are the average of six independent experiments with error bars representing standard deviation. Kd values of each protein are reported in Table 1 as well as within each graph.

We finally proceeded to assess in real-time the RNA unwinding activity of the proteins using a fluorescence-based helicase assay [32]. In this assay, we used a 5’-end Cy5-labeled 5’ovh_dsRNA_26/14_ with a IBRQ quenched at the 3’-end (see Materials and Methods). The unwinding activity was monitored by measuring the increase in fluorescence within the reaction mixture, due to dissociation of the Cy5-labelled RNA strand, as the proteins were incubated in the presence of ATP and Mg^2+^ over a period of 30 minutes (Figure 4). In this experimental condition, E2 displayed the highest RNA unwinding activity (with a rate of 437 fmol of RNA unwound per min, Figure 4C) while E1 an activity 14-fold lower (at 32 fmol of RNA unwound per min, Figure 4A). Surprisingly, the CTE truncations had a dramatic impact on the unwinding activity. E1^1-392^ and E1^1-379^ showed no unwinding capacity (Figure 4B and Figure S5C), and E2^1-384^ retained only 2.4% of the activity observed in the full-length E2 (Figure 4D). In contrast, the R1C2 chimera exhibited an impressive gain in RNA unwinding activity (>300% as compared to E1 and 23% of E2 activity, Figure 4E), while R2C1 displayed activity that was only 5% of E2 and within the range of E1 (∼70%, Figure 4F). Taken together, these results clearly indicate that the CTEs play an essential role in the unwinding activity of the proteins (see Discussion).

**Figure 4.**
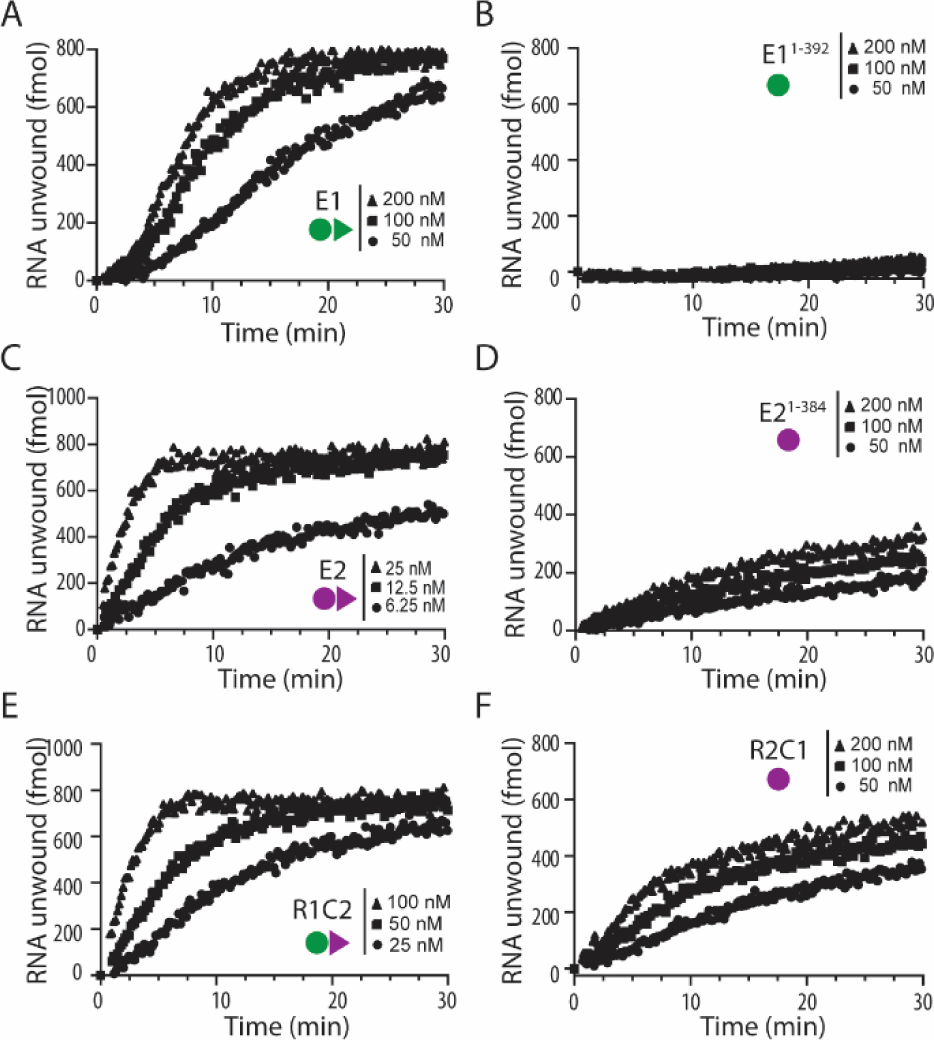
RNA unwinding activity of different RhlE proteins and variants. Reaction mixtures (15 µl) containing 50 mM HEPES (pH 7.2-7.5), 1 mM DTT, 50 mM KGlu, 2 mM MgGlu, 1 mM [γ-32P] ATP, 5 µM duplex RNA and either **(A)** RhlE1 (E1), **(B)** RhlE1^1-392^ (E1^1-392^), **(C)** RhlE2 (E2), **(D)** RhlE2^1-384^ (E2^1-384^), **(E)** R1C2 or **(F)** R2C1 chimera were incubated at 37 °C. The extend of RNA unwound was determined as described in Materials and Methods and was plotted as a function of time. Protein concentrations are specified on each panel. The unwinding rates (min^-1^) for each protein were estimated from the slopes of the curves within the linear range and it is reported in Table 1.

Of note, in ATPase, FP and unwinding experiments the same RNA substrate was used (Figure 2-4). Given our earlier observations of distinct ATPase activities of E1 and E2, depending on the length and structure of the RNA [25], we conducted supplementary ATPase and RNA binding assays using poly (U) RNA and a 31-mer RNA molecule containing an hairpin (Figures S6 and S7, respectively). Similar results were obtained, confirming the consistent effect of the CTEs in these processes.

### RhlE1 and RhlE2 undergo phase separation *in vitro* and form biomolecular condensates *in vivo*

In many instances, IDRs favour the process of liquid-liquid phase separation (LLPS), primarily driven by their weak, multivalent, and transient interactions [13]. Therefore, we conducted an analysis of the phase separation behaviour of RhlE1 and RhlE2, and assessed the influence of the CTEs on LLPS [33]. At concentration of 2.5 µM, both E1 and E2 proteins form droplets *in vitro* in the presence of RNA and the addition of NaCl at a final concentration of 1M resulted in the dissolution of the droplets (Figure 5A). When comparing the two proteins, E2 displays a higher propensity for phase separation than E1. More specifically, E2 forms noticeable droplets at lower concentrations than E1 (0.63 µM vs 1.25 µM, respectively; Figure 5B) and, at similar concentrations, E2 droplets were significantly larger than E1 droplets (Figure 5A mid-panel). Moreover, E2 condensates were less sensitive to salt, forming small droplets in the presence of 250 mM NaCl while E1 droplets formation is impaired at 200 mM NaCl (Figure 5C and Figure S8 A and B).

**Figure 5.**
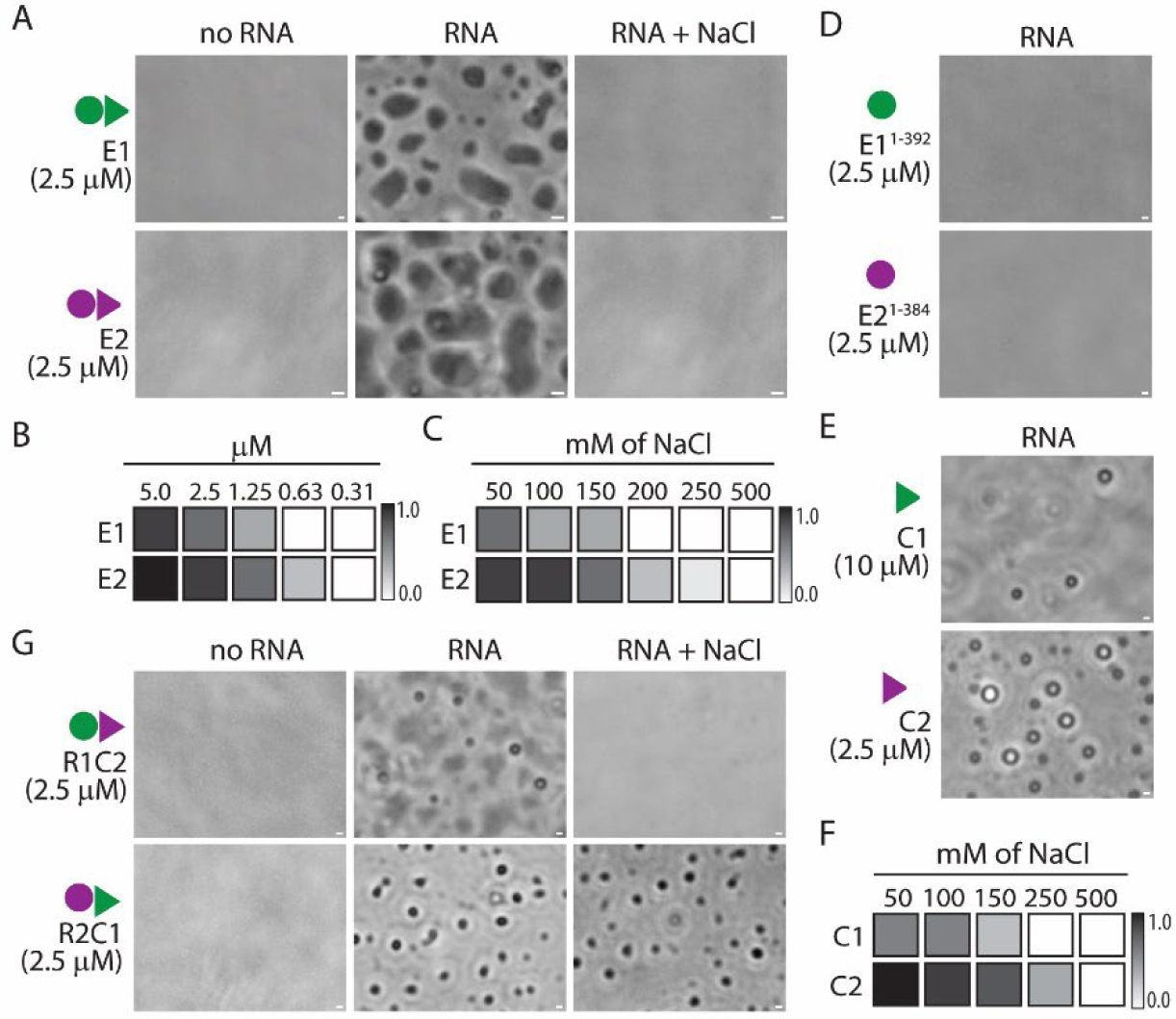
Liquid-liquid phase separation properties of RhlE proteins and their variants. **(A)** In vitro droplet formation of 2.5 µM RhlE1 (E1) or RhlE2 (E2) in the absence (first column) or presence (second column) of 250 ng/µL poly(U) RNA. Images were taken after 60 minutes in presence of absence of RNA. After observing droplet formation, NaCl was added to the reaction mixture at a final concentration of 1M and incubation was continued for 30 minutes (third column). **(B-C)** Droplets formation by E1 and E2 as a function of protein concentration (panel B) and NaCl concentration (panel C) in the mixture. Gray intensities represent the number of droplets in the field, which were normalized for comparison purposes. The maximal droplet number was set to 1, and no droplet formation was set to 0. The raw images are shown in Figure S8 A and B. **(D-E and G)** Droplets formation assay of RhlE1^1-392^ (E1^1-392^) and RhlE2^1-384^ (E2^1-384^) (panel D), RhlE1^379-449^ (C1) and RhlE2^384-634^ (C2) CTE (panel E), R1C2 and R2C1 chimera (panel G). **(F)** Salt sensitivity of C1 and C2 droplets formation. Corresponding raw pictures are reported in Figure S8 C and D. Protein concentrations are indicated in the figure. Each assay was repeated at least three times, and representative images are shown. Scale bars: 5 µm. Of note, all proteins are His10x-Smt3-tagged, and we have checked by Ulp1 cleavage that the tag does not affect the phase separation of the proteins (data not shown).

To assess if RhlE proteins were able to form biomolecular condensates *in vivo*, we expressed msfGFP-tagged RhlE1 or RhlE2 at a similar level in Δ*rhlE1* and Δ*rhlE2* mutant strains, respectively. Both RhlE1- and RhlE2-msfGFP localize into subcellular clusters while the expression of msfGFP alone creates a uniform signal diffused throughout the cytoplasm, suggesting that they could form localized microcompartments (Figure 6A). Chromosomally tagged strains were also tested, but RhlE1-msfGFP signal was barely detectable, probably due to the low expression levels of the protein, while RhlE2-msfGFP clusters were detected (data not shown). The RhlE1- and RhlE2-msfGFP tagged proteins functionality was verified *in vivo* by testing their capacity to recover growth on cold of the Δ*rhlE1* and Δ*rhlE2* mutants (Figure 6B). The purified msfGFP-tagged proteins also formed droplets *in vitro* in the presence of RNA, indicating that the msfGFP-tag did not affect phase separation (Figure 6C). The dynamics of the droplets were investigated by fluorescence recovery after photobleaching (FRAP) experiments (Figure S9). Altogether, the FRAP analysis indicated that RhlE1- and RhlE2-msfGFP droplets had similar recovery half-time (23.9 s and 24.6 s, respectively) but a significantly different mobile fraction (64% and 83%, respectively), which indicates that, although RhlE1 and RhlE2 have similar rates of movement within the droplets (as indicated by the half-time), a higher proportion of RhlE2 is motile as compared to RhlE1.

**Figure 6.**
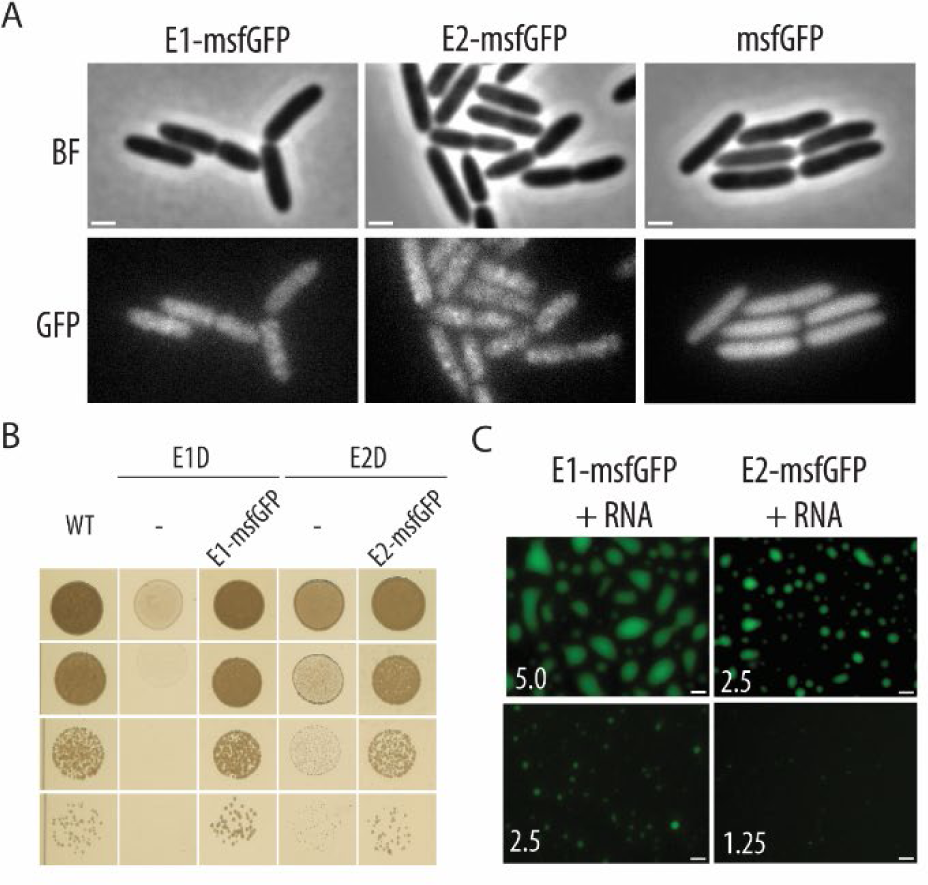
Subcellular localization and phase separation properties of msfGFP-tagged RhlE proteins. **(A)** Localization of RhlE1-msfGFP, RhlE2-msfGFP or msfGFP (green) in *P. aeruginosa* Δ*rhlE1* mutant expressing RhlE1-msfGFP tagged (E1-msfGFP), Δ*rhlE2* mutant expressing RhlE2-msfGFP tagged (E2-msfGFP) or wild type expressing msfGFP only under the control of an AraC-P_BAD_ promoter using mini-Tn7 constructs (see Table S1). Protein expression was induced by growing cells in LB medium with 0.2% of arabinose for two hours. Scale bar: 1 µm, BF: brightfield **(B)** Cold (16 °C) growth of wild type PAO1 (WT), Δ*rhlE1* mutant (E1D), Δ*rhlE1* mutant (E1D) expressing E1-msfGFP, Δ*rhlE2* mutant (E1D) or Δ*rhlE2* mutant (E2D) expressing E2-msfGFP in LB plates with 0.2% arabinose. **(C)** Droplets formation of purified RhlE1-msfGFP (at final concentration of 2.5 or 5 µM) and RhlE2-msfGFP (at final concentration of 1.25 or 2.5 µM) in presence of 250 ng/µl poly(U) RNA. Images were taken after an incubation of 60 minutes. All experiments were performed in independent triplicates and representative results are shown. SDS-page of the purified proteins are shown in Figure S4C.

### The IDRs of RhlE1 and RhlE2 influence phase separation with different strength

Truncation of RhlE1 (E1^1-379^ and E1^1-392^) or RhlE2 (E2^1-384^) prevent the proteins to phase separate in vitro at 2.5 µM, showing that the IDRs are necessary to drive efficient phase separation (Figure 5D), likely through a charge-mediated mechanism [34]. Subsequently, we performed *in vitro* droplet formation assays with the R1C2 and R2C1 chimeras (at 2.5 µM) and we observed that they could both phase separate in the presence of RNA. Nonetheless, droplets formed by the chimeras were smaller than E1 and E2. Moreover, the addition of NaCl at a final concentration of 1M did not result in the dissolution of the R1C2 droplets contrary to what observed with E1 and E2. This could indicate that the droplets formed by R2C1 are of gel-like or solid nature [33].

Importantly, the RhlE1 and RhlE2 CTEs alone (C1 and C2, respectively) could undergo phase separation in presence of RNA (Figure 5E). Similar to E2, C2 formed droplets at a lower concentration than C1 (2.5 versus 10 µM, respectively) and exhibited the capacity to form droplets at higher salt concentrations (Figure 5F). This provides strong evidence that different IDRs confer distinct phase separation properties to the full-length proteins.

### Transcriptome analysis reveals target recognition capacities driven by the RhlE2 IDR/CTE

Previous transcriptome analysis of *P. aeruginosa* mutants carrying *rhlE1* or *rhlE2* deletion and grown at 37°C showed that the former was not significantly affecting cellular transcripts levels while the latter modified ∼15% of the transcriptome, including virulence genes [25]. Here, we conducted an RNA-sequencing analysis from cells grown at 16°C of stains deleted for *rhlE1*, *rhlE2* or both. On this condition, loss of *rhlE1* affected 247 transcripts (198 downregulated and 49 upregulated, *P*-value< 0.05, fold change |FC| ≥2) while 938 transcripts were differentially expressed (209 downregulated and 729 upregulated, *P*-value< 0.05, fold change |FC| ≥2) in the *rhlE2* mutant (Figure 7A and B). Little overlap was observed between RhlE1 and RhlE2 targets, involving in total 77 genes, which confirmed the non-redundancy of the proteins. Of these common targets, 52% were consistently downregulated in both mutants, while 48% showed opposite regulation patterns (Figure 7C). The top KEGG categories overrepresented among the upregulated genes in the E1D strain as compared to the wild type (WT) were related to biofilm formation and alginate biosynthesis, while the most down-regulated functions were sulphur metabolism, active membrane transport, heme, iron starvation and siderophore biosynthesis (Table S4), indicating that the growth defect of the *rhlE1* mutant is probably related to iron limitation [35]. The transcripts affected by the loss of RhlE2 represented instead ∼16% of total cellular transcripts (938/5700), confirming the role of RhlE2 as a global regulator of gene expression. Among the overrepresented upregulated functions in the E2D strain (as compared to the wild type) were genes associated with xenobiotic biodegradation and metabolism, such as 4-fluorobenzoate, chlorocyclohexane and chlorobenzene degradation. Conversely, certain virulence genes like those encoding the type 3 secretion system or T3SS [36] and type 4 pili [37], along with genes involved in aerobic respiration, carbon and nitrogen metabolism, were downregulated (Table S4). Many of these functions were also affected by RhlE2 in the previous RNA-sequencing analysis performed in swarming cells at 37°C [25]. A significant correlation (R=0.552, p<2e-16) was indeed found when comparing the two datasets (log2FC E2D/WT cold *versus* log2FC E2D/WT swarming), despite the cells being subjected to very different growing conditions, indicating that RhlE2 exhibits target specificity (Figure S10).

**Figure 7.**
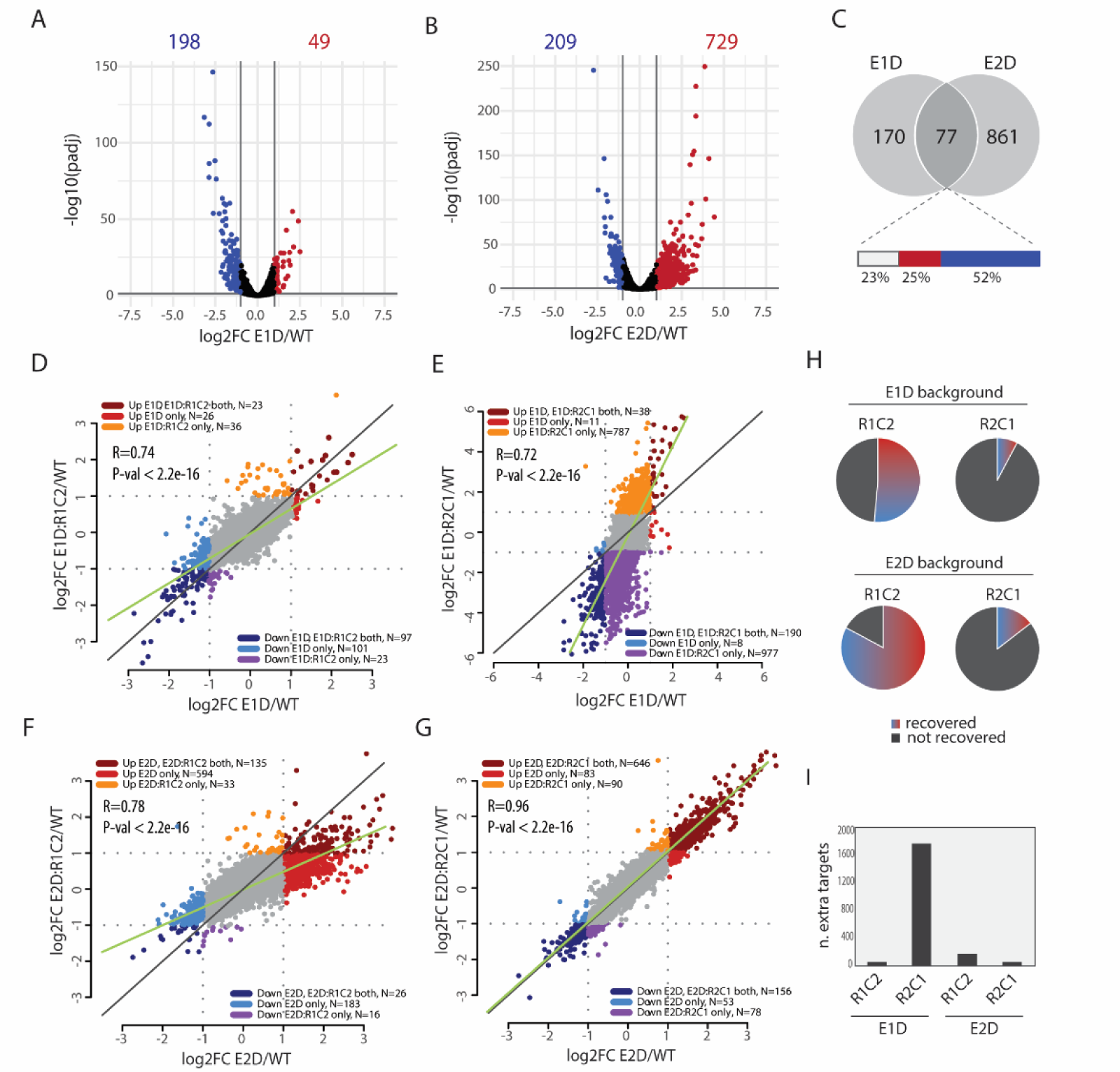
Transcriptome changes associated to RhlE proteins expression in the single *rhlE1* or *rhlE2* mutants. **(A-B)** Volcano plot representation of transcriptome comparison between the wild type PAO1 (WT) and Δ*rhlE1* (E1D, panel A) or Δ*rhlE2* (E2D, panel B) strain. Transcriptome analysis was performed in triplicates, cells from 6 independent spots were grown for 36 hours in LB agar plates at 16°C and pooled together for RNA extraction. The genes are coloured if they pass the thresholds for adjusted P-value (adjp < 0.05) and log fold change |FC| ≥1 (red, upregulated; blue, downregulated). **(C)** Venn diagram illustrating genes commonly regulated by *rhlE1* and *rhlE2* deletion (n=77). The bar represents commonly upregulated (25%, red), commonly downregulated (52%, blue) or oppositely regulated (23%, white) genes. **(D-E)** Scatterplots analysis comparing differentially expressed genes in E1D strain versus E1D strain expressing the R1C2 (panel D) or the R2C1 (panel E) relative to the wild type. **(F-G)** Scatterplots analysis comparing differentially expressed genes in E2D strain versus E2D strain expressing the R1C2 (panel G) or the R2C1 (panel H) relative to the wild type. **(H)** Proportion of “recovered genes” by expression of R1C2 or R2C1 in the E1D or E2D background (see text for more details). **(I)** Number of “extra targets” due to expression of R1C2 or R2C1 in the E1D or E2D background (see text for more details).

Since the targets of RhlE1 and RhlE2 were largely distinct, we hypothesized that analysing the transcriptome of strains expressing R1C2 or R2C1 could aid in distinguishing the contributions of the CTEs to target recognition and, more broadly, in characterising the regulatory action of the chimeras. Scatter plots comparing significantly differentially expressed genes in the E1D strain and the E1D:R1C2 or E1D:R2C1 strains (relative to the wild type), were generated (Figure 7 D and E). The same was performed in the E2D mutant background (Figure 7 F and G). Genes were categorised into six groups, as indicated by different colours in the scatterplots. These categories were based on whether genes were uniquely dysregulated (either up or down, |log2FC| ≥ 1) in one of the two strains, or both. Genes that were uniquely dysregulated in the parental strains (E1D or E2D) and not in the strains expressing the chimera were considered as genes regulated by the chimera, *i.e.,* “recovered genes” (Figure 7H). Vice versa, genes that were uniquely dysregulated in the strains expressing the chimeras, but not in the parental strains (E1D or E2D) were categorised as “extra targets”, *i.e.,* genes not originally regulated by RhlE1 or RhlE2 (Figure 7I). To our surprise, 51% of RhlE1 targets (127 out of 247) and 83% of RhlE2 targets (777 out of 938) were not classified as differentially regulated in the corresponding deleted strain expressing the R1C2 chimera as compared to the wild type, indicating that R1C2 retained some functionality from both RhlE1 and RhlE2 (Figure 7 D and F). In contrast, the R2C1 chimera exhibited very limited RhlE1- or RhlE2-like regulation capacity (19 out of 247 RhlE1 and 136 out of 938 RhlE2 targets recovered) but a significantly higher number of additional targets (1796 genes) in the E1D background, which was not observed in the E2D background (Figure 7E, G and I). The observed growth inhibition of strains expressing the R2C1 chimera in the absence of *rhlE1* (Figure 1C) could be linked to these extra targets.

A similar analysis was also performed in the E1DE2D background to confirm observations made in the single mutants (Figure 8). In total, 906 transcripts were differentially regulated in the E1DE2D mutant as compared to the wild-type: 221 downregulated and 685 upregulated (*P*-value< 0.01, FC>2, Figure 8A). Of these, 120 (92+28) genes out of 221 (54%) and 597 (578+19) out of 685 (87%) were in common with the E2D strain (Figure 8B). In agreement, many of the KEGG functions differentially regulated in E1DE2D were in common with E2D, except the functions regulated by RhlE1, like siderophore biosynthesis, protein export and ribosome (Table S4). Altogether, these data suggest that the gene expression profile of the E1DE2D strain resembles to the sum of transcriptome changes in the E1D and E2D genetic background, reinforcing the non-redundancy model. The R1C2 and R2C1 chimeras had similar effects in the E1DE2D background, with the R1C2 chimera recovering 561 RhlE1 and RhlE2 targets (147+414, Figure 8C), and the R2C1 chimera affecting 743 extra targets (246+497, Figure 8D).

**Figure 8.**
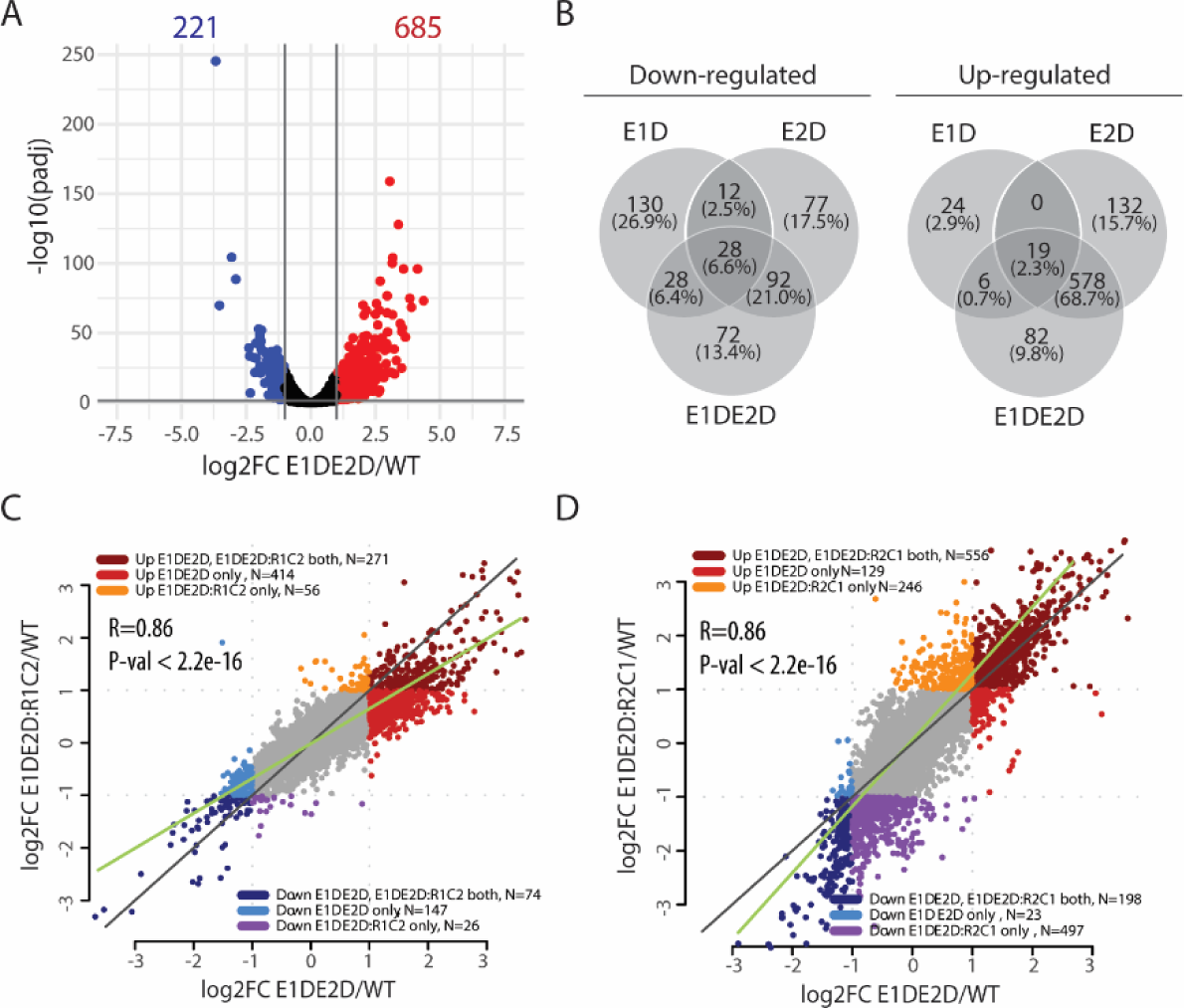
Transcriptome changes associated to expression of RhlE proteins in the Δ*rhlE1*Δ*rhlE2* mutant. **(A)** Volcano plot representation of transcriptome comparison between the wild type PAO1 (WT) and Δ*rhlE1*Δ*rhlE2* (E1DE2D) strain. Transcriptome analysis was performed in triplicates, as explained in Figure 7 legend for single mutants. The genes are coloured if they pass the thresholds for −log10 adjusted P-value (adjp < 0.05) and log fold change |FC| ≥2 (red, upregulated; blue, downregulated). **(B)** Venn diagram illustrating genes commonly regulated by single and double *rhlE1* and *rhlE2* deletion. **(C-D)** Scatterplots analysis comparing differentially expressed genes in E1DE2D strain versus E1DE2D strain expressing the R1C2 (panel C) or the R2C1 (panel D) relative to the wild type.

Overall, we conclude that the R1C2 chimera regulates a subset of both RhlE1 and RhlE2 targets, while the R2C1 chimera has a regulatory activity that differs from either native protein.

### RhlE2 CTE/IDR is sufficient to guide the interaction with the RNase E endonuclease

When comparing the overall transcriptome profiles of all strains through hierarchical clustering, it was intriguing to observe that the strains segregated into two groups based on the presence of C2 (Figure S11A). This observation suggests that the expression of C2 has a global impact on gene expression. Given our previous demonstration that RhlE2 interacts with RNase E via its CTE [25], we hypothesize that this interaction with RNase E and the degradosome underlies the effect of C2. If this hypothesis holds true, then it is conceivable that R1C2 also interacts with RNase E. To test this hypothesis, we conducted bacterial two-hybrid assays, which assess protein interactions based on the proximity of split T25-T18 adenylyl cyclase domains [38]. Consistent with our previous study, co-expression of T25-E2 and T18-RNase E restored the enzyme activity and resulted in a significant increase of β-galactosidase activity as compared to controls. We also observed an interaction between T25-R1C2 and T18-RNase E, but not when using T25-R2C1 nor T25-E1 (Figure 9A). These results confirm our initial hypothesis that C2 is both necessary and sufficient for guiding protein interactions with RNase E.

**Figure 9.**
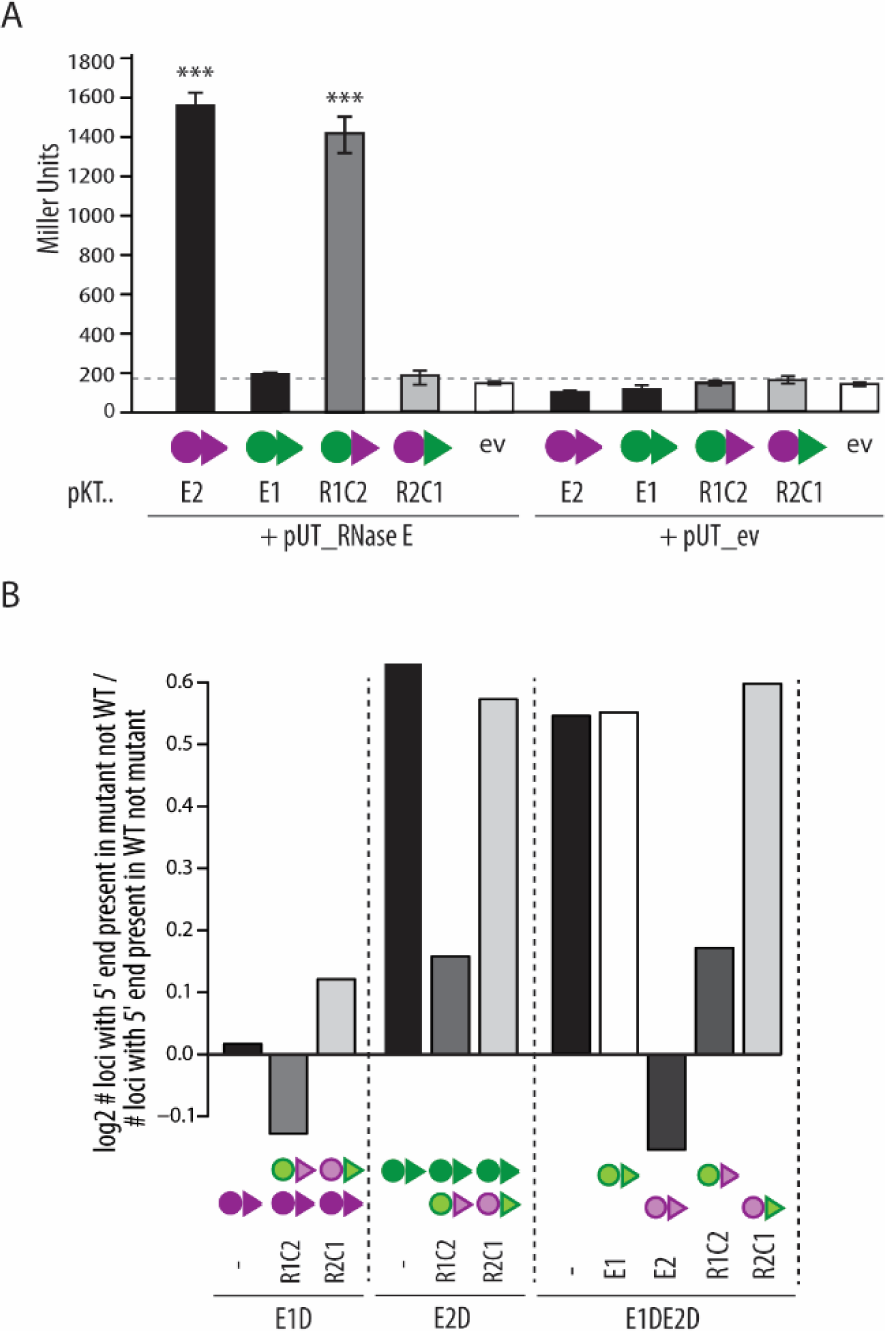
RhlE2 CTE-mediated interaction with RNase E and its effect on RNA decay. **(A)** Reconstitution of adenylate cyclase in the *E. coli* strain BTH101 using a bacterial two-hybrid approach was detected by β-galactosidase assays of lysates of colonies grown on McConkey plates containing 1% maltose, 0.5 mM IPTG, 100 µg/ml ampicillin, and 50 µg/ml chloramphenicol agar plates. Each value is the average of three different cultures ± standard deviation (***, p < 0.01). **(B)** Proportion of loci with 5’ read ends counted only in a mutant strain and not in the wild type (WT) relative to the 5’ read ends (sequenced fragments) counted only in the WT and not in the mutant. A total of 6 million reads was analysed in each strain to exclude an effect due to the library depth (see Materials and methods). Average of the values obtained in the three replicate per strain is plotted while individual values per each replicate is shown in Figure S11B.

In a recent study, de Araújo and colleagues observed an increase of 5’ read ends in the transcriptome of the *rhlE2* mutant as compared to the wild type, suggesting an accumulation of RNA degradation intermediates and an altered RNA decay [29]. We employed the same analysis to assess the efficiency of RNA processing in our strains. Briefly, for each mutant transcriptome, we calculated the ratio between the number of loci with 5’ end present only in the mutant and not in the wild type over the number of loci with 5’ end present only in the wild type and not in the mutant (Figure 9B and S11B). Even though the sequencing protocol produces random read fragmentation and create a background, a specific effect on the number of 5’ ends due to RhlE2 and R1C2 expression was observed. Consistent with the previous observation, deletion of *rhlE2* (E2D or E1E2D strain) led to an increase in 5’ ends compared to the wild type, while the deletion *rhlE1* (E1D) did not. Moreover, the expression of E2, and not E1, significantly decreased the number of 5’ ends (assessed in the E1DE2D background). Interestingly, R1C2 expression in the E1D, E2D, and E1DE2D background also resulted in a decrease in the number of 5’ ends, albeit less pronounced than E2, while no impact was observed for R2C1 expression. Altogether, the analysis reinforced our hypothesis of a role of E2 and R1C2 in RNA processing linked to their interaction with RNase E.

## Discussion

In the context of RNA metabolism, RNA helicases perform diverse and specific functions, often with one RNA helicase unable to substitute for another [25, 39-41]. The specificity of RNA helicases is believed to be conferred by auxiliary domains present in addition to the REC catalytic core that can mediate interactions with RNA targets, protein partners or promote protein dimerization [9-11]. Some RNA-binding auxiliary domains possess target specificity, such as *E. coli* DpbA and *B. subtilis* YxiN CTE, which bind to the loop region of hairpin 92 of the 23S rRNA [42, 43]. Conversely, other RNA-binding auxiliary domains demonstrate broad RNA-binding capabilities and mild selectivity, such as a preference for structured over single-stranded RNAs [9, 11]. Our study focuses on the role of IDRs as ‘auxiliary domains’ of RNA helicases. Interestingly, bacterial RhlE RNA helicases do not possess other structured domains besides the REC core, but a fast-evolving IDR at their C-terminus of different length and amino acid composition (Figure S1).

The RhlE1 and RhlE2 RNA helicases from *P. aeruginosa* are ideal to study both the function and diversity of RhlE IDRs, due to their different CTE/IDR and their specific, non-redundant regulatory functions [25, 41]. Thus, they offer a new frontier for understanding IDR evolution and impact on protein function [44, 45].

We assessed the contribution of the CTEs/IDRs of RhlE1 and RhlE2 to the biochemical activities of the proteins by assessing RNA binding affinity, RNA unwinding and RNA-dependent ATPase activity of protein truncations (missing the CTE) and comparing their biochemical properties to the full-length proteins. In addition, we characterized REC_RhlE1_-CTE_RhlE2_ (R1C2) and REC_RhlE2_-CTE_RhlE1_ (R2C1) chimera to determine the inter-dependency between REC-CTE modules. First, we demonstrate that the CTE of RhlE1 (C1) and RhlE2 (C2) exhibit a strong affinity for 5’ovh_dsRNA26/14 duplex RNA substrate in a 50 mM KGlu environment, with an observed dissociation constants (Kd) of 2.2 nM and 1.3 nM, respectively (Figure 3). Both full-length RhlE1 and RhlE2 demonstrated similar binding affinity to the RNA duplex, with a Kd of 1.7 nM and Kd < 0.8 nM, respectively. The overestimation of the Kd for RhlE2 results from the impracticality of working with RNA concentrations below 0.5 nM due to the detection limit of the plate reader, preventing operating within a binding regime as described by Jarmoskaite et al. (2020) and therefore leading to a definite overestimation of the Kd [46]. Despite the technical limitation, we could show that CTE deletion significantly affects RNA binding in both proteins, as the REC core of RhlE1 (R1) and REC core of RhlE2 (R2) binding to the duplex RNA less efficiently than their full-length protein (43-fold and at least 10-fold decrease, respectively). This finding indicates that, although the REC RNA helicase core possesses RNA-binding characteristics, the RhlE CTEs are crucial for binding RNA with high affinity. Similar results were obtained using a different RNA hairpin substrate, reinforcing our conclusion. Moreover, both R1C2 and R2C1 chimeras exhibited Kd values for 5’ovh_dsRNA26/14 in the same range as the native proteins (1.1 nM and 1.5 nM, respectively). Altogether, our results show that the RhlE1 and RhlE2 CTEs/IDRs can both effectively and similarly increase the RNA binding affinity of RNA helicases and are interchangeable for this function.

A different conclusion is made when observing RNA unwinding activity. Concerning RhlE1, assessing the impact of the CTE on RNA unwinding proved to be challenging as its RNA unwinding activity is inherently weak (an estimation of 0.0317 min^-1^) and observed in reaction mixtures containing an equal amount of RhlE1 and RNA substrate (*i.e.* 50 nM, see Figure 4). The RNA unwinding activity of the R1 is 10-fold lower than RhlE1. Considering that the Kd of the R1 for 5’ovh_dsRNA26/14 is 73 nM, this result strongly suggests that the observed decrease in R1 unwinding activity is primarily due to the decreased binding of RhlE1 to the duplex RNA. Unlike RhlE1, the RhlE2 RNA unwinding activity was measured with an excess of RNA substrate over protein (50 nM RNA versus 6.25 nM of protein) and estimated at ∼0.44 min^-1^. The R2 RNA unwinding activity is 40-fold lower than that of RhlE2, but the unwinding assay was still performed in an RNA substrate concentration higher than the Kd, suggesting that the C2 is also important for the RNA unwinding. The characterization of the chimera activity (R1C2 and R2C1) confirmed our hypothesis of a different contribution of the CTEs to RNA unwinding. Notably, the Kd of both chimeras for 5’ovh_dsRNA26/14 fall within the same nanomolar range of the full-length proteins (Table 1), but the R1C2 chimera exhibits more than threefold increased RNA unwinding activity compared to wild-type RhlE1, suggesting that attaching the C2 to a different REC helicase core can enhance RNA unwinding of a chimera. To our knowledge, this marks the first instance of a chimera demonstrating a gain in activity over a ‘wild-type’ bacterial DEAD-box RNA helicase.

When we tested the RNA-dependent ATPase activity, we worked at saturating RNA concentration (5 µM) for every protein. We could show that the R1 is mildly affected by the CTE (RhlE1^1-392^ = 17.6 min^-1^ versus RhE1 = 18.2 min^-1^). In contrast, the ATPase activity of the R2 is not influenced by the CTE (R2 = 78.5 min^-1^, RhlE2 = 60.9 min^-1^). When comparing the ATPase and RNA unwinding activities of RhlE1 and RhlE2, our finding indicates that these two activities are not always interdependent. Indeed, RhlE1 and RhlE2 have very similar ATPase activities, albeit different RNA unwinding capacities. It has been proposed that DEAD-box helicases frequently undergo futile ATP hydrolysis cycles without dissociating RNA duplexes [47, 48]. We therefore propose that the C2 enhances the release of RNA without affecting ATPase activity.

The difference in the degree of RNA unwinding stimulation between RhlE CTEs may relate to their diverse ability to stimulate phase separation. This has recently been shown for the sexually dimorphic RNA helicase homologs DDX3X and DDX3Y, which possess different IDRs that differentially stimulate the ATPase activity of the protein [49]. Our results indicate that, while RNA induces phase separation of both RhlE1 and RhlE2, RhlE2 exhibits a stronger phase separation propensity than RhlE1. Specifically, RhlE2 phase separation occurs with a lower protein requirement (at least 0.63 µM) and resists higher NaCl concentration (250 mM), while RhlE1 phase separation requires higher protein concentration (a minimum of 1.25 µM) and is more sensitive to NaCl (droplets formation occurs at a concentration below 150 mM). The IDRs alone are necessary and sufficient for phase separation (Figure 5E), and they exhibit the same differences in phase separation propensity and salt sensitivity as the full-length proteins. Altogether, these findings suggest that the IDRs may control RhlE RNA unwinding activity by modifying their localization and concentration within the cell. Fluorescent tagging RhlE1 and RhlE1 *in vivo* confirms that both proteins form clusters within the cell cytoplasm. In eukaryotes, formation of biomolecular condensates via LLPS underpins the biogenesis of a wide array of dynamic membraneless organelles, like the processing bodies, stress granules or Cajal bodies, that are crucial for cell biology and development [50, 51]. Although less characterized in bacteria, biomolecular condensate formation has been recently associated to RNA decay, transcription, or cell division [52]. Future research will examine RhlE1 and RhlE1 localization dynamics under various environmental conditions and the RNA content of these clusters. In addition, it would be interesting to measure the effect of RhlE1 and RhlE2 ATPase activity on the formation of these clusters, since DEAD-box ATP hydrolysis has been shown to trigger the remodelling and disassembly of ribonucleoprotein condensates in both eukaryotes and prokaryotes [21, 53, 54]. Our study also shed some light on the evolution of RhlE RNA helicases. The chimeras do not restore the growth of *rhlE1* and/or *rhlE2* deleted strains at 16°C to wild-type levels, confirming that despite the modular appearance of DEAD-box helicases, core and auxiliary domains are co-dependent for the protein function *in vivo* [38, 39, 46]. To our knowledge, none of the chimeras described to date fully replace *in vivo* the native protein, even if the substrate specificity of certain DEAD-box proteins can be transferred by fusing their specific RNA-binding accessory domain to a different REC core. For instance, the YxiN RNA helicase of *Bacillus subtilis* possesses a CTE auxiliary domain that binds specifically to a fragment (hairpin 92) of 23S rRNA [43]. When the CTER of YxiN is attached to the REC core of the *E. coli* DEAD-box protein SrmB, otherwise known for having broad RNA interaction, specificity for the 23 rRNA is conferred [55]. The transcriptome analyses performed with our strains show that the expression of the R1C2 chimera affects some targets of RhlE1 and some of RhlE2, indicating a partial interchangeability of CTEs and REC domains. Importantly, the interaction of the R1C2 chimera with RNase E confirms our previous findings that the interaction requires the CTE of RhlE2 [25]. Additionally, our results show that C2 alone is sufficient for this interaction, potentially explaining the broad transcriptomic impact of C2 expression (Figure 9 and figure S11). A similar observation was made with *Bacillus subtilis* CshA and CshB DEAD-box helicases. In the Gram-positive bacteria *B. subtilis* and *Staphylococcus aureus*, only the CshA RNA helicase, not CshB, is known to be a global regulator interacting, via its C-terminal end, with the RNase Y endonuclease and the 5′->3 ′ exoribonucleases RNase J1 and RNase J2 [56, 57]. The CshB-ChA chimera, which fuses the C-terminal end of CshA with the REC core of CshB, also interacts with RNase Y [57]. RNase Y plays a pivotal role in the RNA degradosomes of *B. subtilis* and *S. aureus*, akin to the role of RNase E in *E. coli* and other Gram-negative bacteria [58, 59]. Since the last 65 residues of CshA are predicted to have low complexity, this raises the intriguing possibility that phylogenetically distant RNA helicases, such as CshA and RhlE2, have convergently evolved to interact with the primary endonuclease of the RNA degradosome via their IDRs. In this regard, considering the role of C2 on stimulating RNA unwinding, it is possible that RNA helicases interacting with the single-stranded endonucleases, like RNase E or RNase Y, have evolved a strong RNA unwinding activity to help RNA degradation. Contrary to R1C2, R2C1 expression becomes deleterious for RNA metabolism at low temperatures in the absence of RhlE1 and induces more transcriptomic changes than RhlE1 alone. In vitro, R2C1 demonstrates greater RNA unwinding activity compared to RhlE1; however, it also forms irreversible aggregates in the presence of RNA that are resistant to dissolution by salt. Currently, it is unclear which of these two properties is pertinent to the observed phenotype *in vivo*. The lack of a negative impact in the presence of the *rhlE1* gene allows us to infer that the R2C1 chimera does not act as a dominant negative and does not interfere with RhlE1 function. This indicates that the REC core of RhlE2 has evolved to avoid functional redundancy with RhlE1. In other words, after the duplication of the *rhlE* gene, a more sophisticated coordination between the REC core and IDRs likely emerged over time through coevolution, to circumvent redundancy conflicts.

In conclusion, our research highlights the role of IDRs in determining the specific functions of RhlE RNA helicases, imparting unique biochemical and biophysical attributes. Considering the effects of *rhlE2* deletion on *P. aeruginosa* virulence[25], the distinct features of C2 highlighted in this study might help the development of novel therapies against *P. aeruginosa*. Targeting IDRs with small molecules presents some challenges due to their dynamic nature. Nonetheless, the recent discovery of specific small molecules and peptides that modulate IDR-containing proteins—some of which are in clinical trials for cancer-treatment—reveals that these difficulties are not insurmountable [60, 61].

## Materials and Methods

### Growth conditions

Bacterial strains and plasmids used in this study are listed in Table S1. Cells were grown Luria Broth (LB) [62] medium, with shaking at 180 rpm and at 37°C. LB agar (LA) was used as a solid medium. When required, antibiotics were added to these media at the following concentrations: 100 µg/ml ampicillin and 10 µg/ml gentamicin for *E. coli*; and 50 µg/ml gentamicin for *P. aeruginosa*.

### Genetic techniques

PCR and plasmid preparation followed established protocols [63]. DNA cloning was performed by using restriction enzymes or Gibson assembly® (NEB), following manufacturer protocols. Details on the engineered plasmids and bacterial strains can be found in the Table S1 and S2. The oligonucleotides employed in the cloning details are detailed in Table S3. *E. coli* DH5α (utilized for cloning) and *P. aeruginosa* were transformed using heat-shock and electroporation methods, respectively [63]. Verification of all plasmids and strains was done through PCR and Sanger sequencing, which was conducted at Microsynth.

### RNA isolation and sequencing

*P. aeruginosa* strains were cultured at 16°C on LA supplemented with 0.2% arabinose and incubated for 96 hours until colonies became visible. Cells were collected and resuspended immediately on RNA protect Bacteria Reagent (QIAGEN) for RNA stabilization. Total RNA was then extracted and purified utilizing the Monarch Total RNA Isolation Kit (New England Biolabs), followed by triple DNase I (Promega) treatment to eliminate any genomic DNA contamination, and subsequently re-purified using a phenol-chloroform extraction method. Potential DNA contaminants were assessed by performing PCR for 40 cycles using primer pairs targeting *rpoD* and *oprF*, and the integrity of the RNA was verified via agarose gel electrophoresis [25]. Triplicate samples for each strain were prepared. Ribosomal RNA was depleted from the samples, and RNA sequencing was conducted by Novogene.

### RNA sequencing analysis

Reads were mapped using Bowtie2 [64] to *Pseudomonas aeruginosa* PAO1 genome assembly ASM676v1 and counts were aggregated using summarize Overlaps in the package GenomicAlignments [65] in R using the ASM676v1 gtf version 2.2 (NCBI accession: GCF_000006765.1). Differential expression was performed using DESeq2 [66] and transcripts were considered differentially expressed between two conditions if the absolute value of log2FC between conditions was greater than 1 (i.e. 2-fold up or down) with a p-value less than 0.05. To find functional enrichment among differentially expressed genes groups, the Kyoto Encyclopedia of Genes and Genomes (KEGG) was used for gene set definitions, downloaded using the package KEGGREST [67] in R. Enrichment was calculated using a hypergeometric test. All correlations presented are Pearson’s product moment correlation coefficient. Analysis of 5’ read ends per loci was performed as described by de Araújo et al. [29] except that each dataset read number was downsampled to 6 million.

### Protein purification

His_10_Smt3-tagged RhlE1 (E1), RhlE1^1-392^ (E1^1-392^), RhlE1^1-379^ (E1^1-379^), RhlE2 (E2), RhlE2^1-384^ (E2^1-384^), REC_RhlE1_-CTE_RhlE2_ (R1C2) and REC_RhlE2_-CTE_RhlE1_ (R2C1), RhlE1^379-449^ (C1), RhlE2^384-639^ (C2), RhlE1-msfGFP (E1-msfGFP) and RhlE2-msfGFP (E2-msfGFP) proteins were purified from soluble bacterial lysates by nickel-agarose affinity chromatography as described previously in the supplementary information of Hausmann *et al.* 2021 [25]. The recombinant His_10_Smt3-tagged proteins were recovered predominantly in the 250 mM imidazole fraction. The imidazole elution profiles were monitored by SDS-PAGE (250 mM imidazole fraction for every recombinant protein is shown in Figure S4). For LLPS experiments (see below), the 250 mM imidazole fraction was concentrated with a centrifugal filter (Amicon Ultra 4; 10’000 MWCO) and then stored at -80°C. The protein concentration was determined using the Bio-Rad dye reagent with BSA as the standard.

### ATPase reaction

Reaction mixtures (15 µl) containing 50 mM HEPES (pH7.2-7.5), 1 mM DTT, 2 mM MgGlu, 1mM ATP + traces of [γ-32P]ATP, 50 mM KGlu, RNA as specified, and enzyme as specified were incubated for 15 min at 37°C. The reactions were quenched by adding 3.8 µl of 5 M formic acid. Aliquots (2 µl) were applied to a polyethylenimine (PEI)-cellulose thin layer chromatography (TLC) plates (Merk), which were developed with 1 M formic acid, 0.5 M LiCl. ^32^Pi release was quantitated by scanning the chromatogram with laser Scanner Typhoon FLA 7000 (General Electric).

### RNA binding assay

RNA binding was measured by fluorescence polarization assay (FP) as described elsewhere [26, 68, 69]. RNA was purchased from IDT (LubioScience GmbH, Zurich, Switzerland). Assays were performed in 384-well black plates (Corning 3820). Reaction mixtures (20 µl) containing 50 mM HEPES (pH 7.2-7.5), 5 mM DTT, 2 mM MgGlu, 1 mM non-hydrolysable ATP analogue adenosin-5′-(β,γ-imido)triphosphate (AMP-PNP), 50 mM KGlu, 0.5 nM 5’-FAM-labeled duplex RNA (5’ovh_dsRNA_26/14_; 5’-FAM-AACAAAACAAAA<u>UAGCACCGUAAAGC</u>/ 5’-<u>GCUUUACGGUGCUA</u>), or 0.5 nM 5’-FAM-labeled 31-mer (5’-FAM-CAGGUCCCAAGGGUUGGGCUGUUCGCCCAUU; this oligonucleotide contains the stem of hairpin 92 of 23S rRNA [68], and proteins concentration as specified were incubated at 25 °C for 15 min. Fluorescence anisotropy measurements were performed using a Victor Nivo plate reader (PerkinElmer, Basel, Switzerland). The excitation wavelength was set to 480 nM and the emission was measured at 530 nm. Data points were analysed as a set of triplicates for each sample. The measured anisotropy was normalized in comparison to the protein-free sample. Values represent the means of six independent experiments +/- standard deviation. To extract binding constants, changes in fluorescence polarization were plotted against protein concentration and the data were fitted to a single exponential Hill function [70] using GraphPad Prism (GraphPad Software, Inc.). The Kd of RhlE2 full length is very close to the RNA concentration employed in the assay, suggesting that we are not within the ‘binding regime’ [46]. Since the fluorescence detection limit impairs the utilization of RNA concentrations below 0.5 nM, the overestimated Kd values are denoted by the ‘<’ sign in the results. Note that fluorescence polarization was found stable when measured within the range of 5 to 20 minute (data not shown). Reaction mixtures used to determine the Kd of RhlE1 CTE (C1) and RhlE2 CTE (C2) do not include the non-hydrolysable ATP analogue adenosin-5′-(β,γ-imido) triphosphate.

### Helicase assay

RNA unwinding was monitored with a fluorescence-based approach using two modified RNA strands as described in Ali R OÖ zes et al. [32]. RNA substrate purchased from IDT contains a 14-base pair duplex region and a 12-nucleotides single strand region 5’overhang consisting of 5’-AACAAAACAAAA<u>UAGCACCGUAAAGC</u>-3’-(IBRQ) / 5’-(Cy5)-<u>GCUUUACGGUGCUA</u> (5’ovh_dsRNA_26/14_). In this assay, an increase of fluorescence is measured when the Cy5 RNA strand was separated from the complementary strand carrying the quencher (IBRQ). This RNA unwinding assay was performed at 37 °C in 384-well plates using a Victor Nivo plate reader (PerkinElmer, Basel, Switzerland). The excitation wavelength was set to 640 nM and the emission was measured at 685 nm. Reaction mixtures (15 μl) containing 50 mM HEPES, 5 mM DTT, 2 mM MgGlu, 1 mM ATP (or no ATP), 50 mM KGlu, 50 nM of 5’ovh_dsRNA_26/14_, 3 μM of RNA competitor (5’-GCUUUACGGUGCUA), 1.2 U/µl RNasin plus (Promega) and purified recombinant protein as specified were incubated at 37 °C. RNA competitor was added in order to prevent the re-annealing of the Cy5 -RNA to 26-mer RNA strand containing the quencher. The fluorescence change was measured over time, starting 1 minute after the addition of the protein. Data from fluorescence measurement were baseline corrected by subtracting the fluorescence of the ‘No helicase’ control included in every experiment. To determine the amount of unwound RNA in (fmol/min), data were normalized in respect to the fluorescence signals for the fully dissociated 5’ovh_dsRNA_26/14_ and the fully quenched (annealed) 5’ovh dsRNA_26/14_, respectively. The maximum fluorescence was assessed in a reaction mixture lacking protein and heated at 90°C for 5 minutes. The recorded maximum fluorescence closely aligns with the value obtained using 70 nM RhlE2 (not shown), which was consistently included as a positive control in each experiment. An estimation of unwinding constant rates of each protein were determined in the linear range of the curve shown in Figure 4 and Figure S5C.

### *In vitro* liquid droplet reconstitution assay (or droplet assembly)

The in vitro liquid-liquid phase separation (LLPS) experiment was conducted at at 25 °C in an Eppendorf tube with a total volume of 25 µl. Proteins were diluted to in buffer H (50 mM HEPES (pH 7.2-7.5), 500 mM NaCl, 0.01 % Triton X-100, and 5µl diluted protein were transferred to the Eppendorf tube containing 20 µl reaction mixtures. These reaction mixtures (25 µl total) contain 50 mM HEPES (pH7.2-7.5), 0.4 µg/µl BSA, 1 mM DTT, 2m MgCl_2_, 100 mM NaCl, 250 ng/µl Poly(U), and enzyme as specified were incubated for 10 min at 25 °C and then transferred into a 384-well plate (Starlab 4TI-0214) for 50 min. The images were taken using a Nikon Ts2R-FL microscope (magnification 40X). Note that for RhlE1-msfGFP, and RhlE2-msfGFP, the images were taken using a EVOS FL (life technologies) microscope (magnification 40X).

### Bacterial two-hybrid assay

Bacterial two hybrid experiments were performed as previously described [25]. Briefly, recombinant pKT25 and pUT18C derivative plasmids were transformed simultaneously into the *E. coli* BTH101 strain. Transformants were spotted onto McConkey agar plates supplemented with 1 mM isopropyl β-d-thiogalactoside (IPTG) in the presence of 100 μg/ml ampicillin, 50 μg/ml kanamycin. Positive interactions were identified as blue colonies after 24h incubation at 30°C and quantified by β-galactosidase assays. Data are mean values of three independent samples ± standard deviations.

### Fluorescence microscopy

Phase contrast and fluorescence microscopy were performed on a Zeiss Axio Imager 2 equipped with an objective alpha Plan-Apochromat 100x/1,46 Oil Ph3 M27 and a ZEISS Axiocam 305 mono CCD camera. Bacteria (30 µl of an overnight culture) were grown for 2 hours on 3 ml of LB with 0.2% arabinose before placing them on a patch consisting of 1% agarose (Sigma) in water. Images were processed with Image J (NIH, USA).

## Data availability

RNA-sequencing raw files and read counts per gene have been deposited in the NCBI Gene Expression Omnibus (GEO) database under the accession number XXXX.

## Acknowledgments.

We are grateful to Nathanaël Guggenheim for the purification of RhlE1- and RhlE2-msfGFP tagged proteins. We also thank Julian Prados (Bioinformatic facility at University of Geneva) for the helpful discussion concerning RNA sequencing data analysis.

