## Supplementary file for "Intrinsically disordered regions regulate RhlE RNA helicase functions in bacteria"

**Content of Supplementary information:**

Supplementary Tables

Supplementary Materials and Methods

Supplementary Figures Legends

Supplementary references

Supplementary Figures

### Supplementary Tables

### Table S1. Strains and plasmids used in this study.

| **Strains** | **Genotype/relevant characteristics** | **Source** |
|---|---|---|
| *E. coli* | | |
| *Rosetta* (DE3) | *F- ompT hsdSB(rB- mB-) gal dcm* (DE3) pRARE (Cm$^R$) | Lab collection |
| DH5α | *recA1 endA1 hsdR17 supE44 thi-1 gyrA96 relA1* Δ(*lacZYA-argF*)*U169* [Φ80d*lacZ*M*15*]F$^-$ NaI$^r$ | [1] |
| HB101 | *proA2 hsdS20*($r_{B^-}$ $m_{B^-}$) *recA13 ara-14 lacYI galK2 rpsL20 supE44 xyl-5 mtl-1* F$^-$ | [1] |
| *P. aeruginosa* | | |
| PAO1 | Wild-type (WT) | [2] |
| E1D | PAO1 containing a 1332-bp deletion in the *rhlE1* (PA3950) locus | [3] |
| E2D | PAO1 containing a 1902-bp deletion in the *rhlE2* (PA0428) locus | [3] |
| E1DE2D | PAO1 with a *rhlE1* and *rhlE2* deletion | [3] |
| E1::msfGFP | PAO1 with a *rhlE1::msfGFP* | This study |
| E2::msfGFP | PAO1 with a *rhlE2::msfGFP* | This study |
| **Plasmids** | | |
| pET28b-10xHis-Smt3 | Broad host range vector for expression of N-terminal 10xHis-Smt3-tag proteins, Km$^r$ | [4] |
| pKT25 | Two-hybrid plasmid, *cyaAT25* fusion, Km$^R$ | [5] |
| pUT18C | Two-hybrid plasmid, *cyaAT18* fusion, Ap$^R$ | [5] |
| pME6182 | Mini-Tn*7* gene delivery, ColE1 replicon; Gm$^r$Ap$^r$ | [6] |
| pLM37 | pET28b-10xHis-Smt3 derivative for His-Smt3_RhlE2 purification, Km$^r$ | [3] |
| pLM38 | pET28b-10xHis-Smt3 derivative for His-Smt3_RhlE1 purification, Km$^r$ | [3] |
| pLM118 | pLM37 derivative with for His-Smt3_REC$_{RhlE1}$CTE$_{RhlE2}$ (R1C2) purification, Km$^r$ | This study |
| pLM119 | pLM37 derivative with for His-Smt3_ REC$_{RhlE2}$CTE$_{RhlE1}$ (R2C1) purification, Km$^r$ | This study |
| pLM173 | pLM37 derivative with for His-Smt3_RhlE1$^{1\text{-}379}$ (R1.0) purification, Km$^r$ | This study |
| pLM172 | pLM37 derivative with for His-Smt3_RhlE1$^{1\text{-}392}$ (R1) purification, Km$^r$ | This study |
| pLM174 | pLM37 derivative with for His-Smt3_RhlE2$^{1\text{-}384}$ (R2) purification, Km$^r$ | This study |
| pLM534 | pLM37 derivative with for His-Smt3_RhlE1-msfGFP (E1-msfGFP) purification, Km$^r$ | This study |
| pLM545 | pLM37 derivative with for His-Smt3_RhlE2-msfGFP (E2-msfGFP) purification, Km$^r$ | This study |
| pLM176 | pLM37 derivative with for His-Smt3_RhlE1$_{379\text{-}449}$ purification, Km$^r$ | This study |
| pLM175 | pLM37 derivative with for His-Smt3_RhlE2$_{384\text{-}634}$ purification, Km$^r$ | This study |
| pLM86 | pKT25 derivative for T25-RhlE1expression, Km$^r$ | [3] |
| pLM85 | pKT25 derivative for T25-RhlE2 expression, Km$^r$ | [3] |
| pLM195 | pKT25 derivative for T25-R1C2 expression, Km$^r$ | This study |
| pLM196 | pKT25 derivative for T25-R2C1 expression, Km$^r$ | This study |
| pLM88 | pME6182 derivative carrying $P_{BAD}$_3XFLAG-RhlE2, Gm$^r$ | [3] |
| pLM128 | pME6182 derivative carrying $P_{BAD}$_3XFLAG-RhlE1; Gm$^r$ | [3] |
| pLM502 | pME6182 derivative carrying $P_{BAD}$_3XFLAG-RhlE1-msfGFP; Gm$^r$ | This study |
| pLM503 | pME6182 derivative carrying $P_{BAD}$_3XFLAG-RhlE2-msfGFP; Gm$^r$ | This study |
| pLM508 | pME6182 derivative carrying $P_{BAD}$_3XFLAG-RhlE1$^{1\text{-}379}$; Gm$^r$ | This study |
| pLM202 | pME6182 derivative carrying $P_{BAD}$_3XFLAG-RhlE1$^{1\text{-}392}$; Gm$^r$ | This study |
| pLM510 | pME6182 derivative carrying $P_{BAD}$_3XFLAG-RhlE2$^{1\text{-}384}$; Gm$^r$ | This study |
| pLM116 | pME6182 derivative carrying $P_{BAD}$_3XFLAG-R1C2; Gm$^r$ | This study |
| pLM117 | pME6182 derivative carrying $P_{BAD}$_3XFLAG-R2C1; Gm$^r$ | This study |
| pLM509 | pME6182 derivative carrying $P_{BAD}$_3XFLAG-RhlE1$^{379\text{-}449}$; Gm$^r$ | This study |
| pLM180 | pME6182 derivative carrying $P_{BAD}$_3XFLAG-RhlE2$^{384\text{-}634}$; Gm$^r$ | This study |
| pLM147 | pUT18C derivative for T18-RNaseE expression, Ap$^r$ | This study |
| pRK2013 | Helper plasmid; Tra$^+$ Km$^r$ | [1] |

**Table S2. Cloning details.**

| Plasmid | Backbone | Insert | Cloning method |
|---|---|---|---|
| pLM118 | pET28b-10xHis-Smt3 | p95-p215 and p216-p96 using PAO1gDNA as template, then p95-p96 overlapping PCR | HindIII, XhoI |
| pLM119 | pET28b-10xHis-Smt3 | p85-p217 and p218-p86 using PAO1gDNA as template, then p85-p86 overlapping PCR | HindIII, XhoI |
| pLM173 | pET28b-10xHis-Smt3 | p95-p348 using PAO1gDNA as template | HindIII, XhoI |
| pLM172 | pET28b-10xHis-Smt3 | p95-p289 using PAO1gDNA as template | HindIII, XhoI |
| pLM174 | pET28b-10xHis-Smt3 | p85-p347 using PAO1gDNA as template | HindIII, XhoI |
| pLM534 | pET28b-10xHis-Smt3 | p95-p605 using E1::msfGFP gDNA as template | HindIII, XhoI |
| pLM545 | pET28b-10xHis-Smt3 | p85-p605 using E2::msfGFP gDNA as template | HindIII, XhoI |
| pLM176 | pET28b-10xHis-Smt3 | p317-p96 using PAO1gDNA as template | HindIII, XhoI |
| pLM175 | pET28b-10xHis-Smt3 | p313-p86 using PAO1gDNA as template | HindIII, XhoI |
| pLM195 | pKT25 | p146-p149 using pLM118 as template | BamHI, EcoRI |
| pLM196 | pKT25 | p148-p147 using pLM119 as template | BamHI, EcoRI |
| pLM502 | p140-p141 with pLM88 as template | p141-p536 using E1::msfGFP gDNA as template | Gibson assembly® |
| pLM503 | p140-p141 with pLM88 as template | p293-p536 using E2::msfGFP gDNA as template | Gibson assembly® |
| pLM508 | p140-p141 with pLM88 as template | p142-p350 using gDNA as template | Gibson assembly® |
| pLM509 | p140-p141 with pLM88 as template | p142-p292 using gDNA as template | Gibson assembly® |
| pLM510 | p140-p141 with pLM88 as template | p293-p349 using gDNA as template | Gibson assembly® |
| pLM116 | p140-p141 with pLM88 as template | p142-p133 using pLM118 as template | Gibson assembly® |
| pLM117 | p140-p141 with pLM88 as template | p132-p143 using pLM119 as template | Gibson assembly® |
| pLM202 | p140-p141 with pLM88 as template | p143-p318 using gDNA as template | Gibson assembly® |
| pLM180 | p140-p141 with pLM88 as template | p531-p149 using gDNA as template | Gibson assembly® |

### Table S3: Primer list

| NAME | SEQUENCE (5'-3') |
|---|---|
| p85 | CAGTAAGCTTCATCCTTTTCTTCCCTCGGACTCTCC |
| p86 | CATGCTCGAGTTAGCGGTTGCGGGTCAGCAG |
| p95 | CAGTAAGCTTCAACGTTCGCTTCCCTCGGTCTGCTC |
| p96 | CATGCTCGAGCTATTTGCCGGACGGCCGCTTC |
| p132 | CGGTTTACAAGCATAAAGCTTCTCGAGGTCGACGGTATCG |
| p133 | GGCAGGCATGCCATGGTACCTCTAGAACTAGTTTAGCGG |
| p140 | CTTGTCATCGTCATCCTTGTAATC |
| p141 | GGTACCATGGCATGCCTG |
| p142 | GATGACGATGACAAGACGTTCGCTTCCCTCGG |
| p143 | CAGGCATGCCATGGTACCCTATTTGCCGGACGGCC |
| p146 | GTACGGATCCCACGTTCGCTTCCCTCGGTC |
| p147 | GATCGAATTCCTATTTGCCGGACGGCCGC |
| p148 | GTACGGATCCCTCCTTTTCTTCCCTCGGACTC |
| p149 | GATCGAATTCTTAGCGGTTGCGGGTCAGC |
| p149 | GATCGAATTCTTAGCGGTTGCGGGTCAGC |
| p215 | GACTTCGAGCCGGAGCACGTGCTGCCCGAGGTGGCC |
| p216 | GGCCACCTCGGGCAGCACGTGCTCCGGCTCGAAGTC |
| p217 | GGTTTCGACCCCGAGGCCCGGGTGCCGCAGACCGCA |
| p218 | TGCGGTCTGCGGCACCCGGGCCTCGGGGTCGAAACC |
| p289 | CATGCTCGAGCTACTTCAGCACCACGCCGCCCG |
| p292 | CAGGCATGCCATGGTACCCTACTTCAGCACCACGCCGCC |
| p293 | GATGACGATGACAAGTCCTTTTCTTCCCTCGGACTC |
| p313 | CAGTAAGCTTCAGGTGGTGGTTCCGGTACCGAGAAGCCGGCTG |
| p317 | CAGTAAGCTTCAGGTGGTGGTTCCAAGAAACCGAAGAAGCCG |
| p318 | GATGACGATGACAAGAAGAAACCGAAGAAGCCG |
| p347 | CATGCTCGAGTTAGGCCTCGGGGTCGAAACC |
| p348 | CATGCTCGAGCTAGTGCTCCGGCTCGAAGTCGG |
| p349 | CAGGCATGCCATGGTACCCTAGGCCTCGGGGTCGAAACC |
| p350 | CAGGCATGCCATGGTACCCTAGTGCTCCGGCTCGAAGTCGG |
| p531 | GTACGGATCCCGCCGTGCTGCCCGAGG |
| p536 | CAGGCATGCCATGGTACCTTATTTGTAGAGTTCATCC |
| p605 | CATGCTCGAGTTATTTGTAGAGTTCATCCATGCCGT |

### Supplementary Materials and methods

**Western blot analysis.** For each strain, cells density was normalized to an OD600 of 1.0 before loading it onto a 4-12% Bis-Tris mPAGE gel. The gel was migrated and transferred to a nitrocellulose membrane at 3 mA/cm2. After transfer, membranes were blocked overnight in blocking buffer (5% milk powder, 0.1% Tween 20 in Tris-buffered saline, pH 8.0). A 1:10000 dilution of primary anti-Flag antibody (Sigma, F3165) was used. Secondary anti-mouse antibodies conjugated to horseradish peroxidase (Sigma, A9044) were used at a dilution of 1:5000. Western blots were developed using Super-Signal West Pico Chemiluminescent Substrate (Pierce) and visualized on a LAS3000 Fuji Imager.

**Fluorescence recovery after photobleaching (FRAP).** The FRAP assays were conducted using the bleaching module of Nikon A1r+ confocal microscope for RhlE1 and RhlE2 droplets individually. Bleaching of msfGFP signal was focused on a circular region of interest (ROI) using 100% 488 nm laser power for 500 ms, and time-lapse images were collected afterward every 5 seconds for 5 minutes. A same-sized circular area away from the bleaching point was selected as an unbleached control. The fluorescence intensity was directly measured in the Zen software and reported as relative to pre-bleaching time points. The halftime for each replicate and mobile fraction was calculated as previously described [7]. Experiments were performed in triplicates and the two-tail t-test was used to calculate the p-values using GraphPad Prism (GraphPad Software, Inc.).

**Supplementary Figures Legends**

**Figure S1: RhlE proteins C-terminal disordered region. (A)** Sequence alignment of *P. aeruginosa* RhlE1 (green) and RhlE2 (violet). Highlight indicated the REC catalytic domain. **(B)** Disordered region prediction of RhlE proteins of *P. aeruginosa* (RhlE1 and RhlE2), *Escherichia coli* ($RhlE_{Ec}$) and *Caulobacter crescentus* ($RhlE_{Cc}$) using IUPRED [8].

**Figure S2: Western blot showing expression of 3xFLAG-tagged RhlE protein variants.** 3xFLAG-tagegd RhlE protein variants were expressed under the control of an AraC-$P_{BAD}$ promoter and cloned into Tn7, which is inserted into the chromosome of the receiver strain Δ*rhlE1* (E1D), Δ*rhlE2* (E2D), or Δ*rhlE1*Δ*rhlE2* (E1DE2D) at the attTn7 site (see Table S1 and S2). Expression of the RhlE proteins and their variants (E1, E2, R1C2, or R2C1) was induced with 0.2% arabinose, cells were collected to an OD600 of 1.0. FLAG detection by western blot was performed using anti-FLAG antibodies (top panel), a commercial anti-RpoB antibody (663905, BioLegend) was used as loading control (bottom panel). See Materials and methods for details.

**Figure S3: Complementation assays. (A-B)** Growth at 16 °C of wild type PAO1, E1D mutant **(A)** or E2D mutant **(B)** expressing 3xFLAG-tagged RhlE REC core (R1 or R2) or CTE (C1 or C2) in LA plates + 0.2% arabinose. **(C)** Growth at 16 °C of wild type PAO1, E1D mutant and E1D mutant expressing 3xFLAG-tagged RhlE1 and the chimeras (R1C2 or R2C1) in LB liquid + 0.2% arabinose. **(D)** Growth at 37 °C of wild type PAO1, Δ*rhlE1* mutant (E1D), Δ*rhlE2* mutant (E2D) and Δ*rhlE1*Δ*rhlE2* mutant (E1DE2D), along with their corresponding strains expressing a 3xFLAG-tagged RhlE variant (either E1, E2 R1C2, or R2C1) under the control of an AraC-$P_{BAD}$ promoter using mini-Tn7 constructs.

**Figure S4: RhlE proteins purification**. **(A-C)** Aliquots (2.5 µg) of the nickel-agarose preparations of His10x-Smt3-tagged RhlE1 (E1), RhlE2 (E2), R1C2 and R2C1 chimeras, and mutant $RhlE1^{1-379}$ ($E1^{1-379}$), $RhlE1^{1-392}$ ($E1^{1-392}$), $RhlE2^{1-384}$ ($E1^{1-384}$), $RhlE1^{379-449}$ (C1), $RhlE2^{384-634}$ (C2), RhlE1-msfGFP (E1-msfGFP), RhlE2-msfGFP (E2-msfGFP), were analysed by SDS-PAGE. The polypeptides were visualized by staining with Coomassie Blue dye. The positions and sizes (kDa) of marker polypeptides are indicated on the left.

**Figure S5: Characterization of $RhlE1^{1-379}$ truncation. (A)** ATPase activity **(B)** RNA binding **(C)** RNA unwinding **(D)** phase separation of $RhlE1^{1-379}$ truncation. Experiments were performed as described in Figure 2-5 legend, respectively, and in the Material and methods.

**Figure S6: RNA-dependent ATPase activity of different RhlE proteins and their variants.** Reaction mixtures (15 µl) containing 50 mM HEPES (pH 7.2-7.5), 1 mM DTT, 50 mM KGlu, 2 mM MgGlu, 1 mM [γ -32P] ATP, 250 ng/µl poly(U) RNA and either **(A)** RhlE1 (E1), **(B)** RhlE1$^{1-379}$ (E1$^{1-379}$), **(C)** RhlE2 (E2), **(D)** RhlE2$^{1-384}$ (E2$^{1-384}$), **(E)** R1C2 or **(F)** R2C1 chimera and **(G)** RhlE1$^{1-392}$ (E1$^{1-392}$) were incubated for 15 min at 37 °C. Pi release was determined as described in Materials and Methods and was plotted as a function of protein concentration. Data are the average of three independent experiments with error bars representing standard deviation. **(H)** Bar plot illustrating the specific ATPase activity for each protein, which was calculated from the slope of the titration curve (shown in panel A to F) in the linear range.

**Figure S7. RNA binding of different RhlE proteins and variants measured by fluorescence polarization assays.** Reaction mixtures (20 µl) containing 50 mM HEPES (pH 7.2-7.5), 1 mM DTT, 50 mM KGlu, 2 mM MgGlu, 1 mM non-hydrolysable ATP analogue, 0.5 nM 5'-FAM-labeled 31-mer RNA [9] and either **(A)** RhlE1 (E1), **(B)** RhlE1$^{1-379}$ (E1$^{1-379}$), **(C)** RhlE2 (E2), **(D)** RhlE2$^{1-384}$ (E2$^{1-384}$), **(E)** R1C2 or **(F)** R2C1 chimera and **(G)** RhlE1$^{1-392}$ (E1$^{1-392}$), **(H)** RhlE1$^{379-449}$ -named C1-, **(I)** RhlE2$^{384-634}$ -named C2- protein as specified were incubated for 15 min at 25 °C. mFP was determined as described in Material and Methods and was plotted as of function of protein concentration. **(J)** Kd values of each protein are reported in the bar plot as well as within each graph. Data are the average of six independent experiments with error bars representing standard deviation.

**Figure S8: Extended figure of phase separation properties of RhlE proteins and variants. (A)** In vitro droplet formation of RhlE1 (E1) or RhlE2 (E2) at different protein concentration (from 5 to 0.31 µM as indicated) and in presence of 250 ng/µL poly(U) RNA. **(B)** Effects of salt concentration on E1 and E2 (both at 2.5 µM) droplets. After observing droplets formation of E1 or E2, NaCl was added to the solution at a final concentration ranging from 50 to 500 mM as indicated. **(C)** Droplets formation by RhlE1$^{379-449}$ (C1) and RhlE2$^{384-634}$ (C2) as a function of protein concentration in the mixture. **(D)** Effect of salt on droplets by C1 (at 10 µM) and C2 (at 2.5 µM) as performed for full-length proteins. Each assay was repeated at least three times, and representative images are shown. Scale bars: 5 µm.

**Figure S9: FRAP experiments. (A)** Time-lapse images of in vitro FRAP experiments. The FRAP experiments were performed identically for RhlE1-msfGFP and RhlE2-msfGFP droplets at 1.75 µM. **(B)** FRAP curves for in vitro droplets of RhlE1-msfGFP (green) and RhlE2-msfGFP (violet). The traces of the FRAP data represent mean ± SEM from three independent experiments. **(C-D)**

Half-time and mobile fractions from panel B. A two-tailed t test was used to calculate the p value. ***p < 0.01.

**Figure S10: The RhlE2 regulon in different growing conditions.** Scatterplot analysis comparing differentially expressed genes in E2D strain swarming or growing on cold (16° C) relative to wild type. Blue dots correspond to virulence or virulence-associated genes according to the Pseudomonas.com database. R= Pearson's product moment correlation coefficient.

**Figure S11: Hierarchical clustering of expression profiles of strains expressing RhlE proteins variants growing on cold.** Strains whose transcriptome profile was analysed were: wild-type (WT), Δ*rhlE1* (E1D), Δ*rhlE2* (E2D), Δ*rhlE1*Δ*rhlE2* (E1DE2D), E1D expressing 3xFLAG tagged R1C2 chimera (E1D:R1C2), E1D expressing 3xFLAG tagged R2C1 chimera (E1D:R2C1), E2D expressing 3xFLAG tagged R1C2 chimera (E2D:R1C2), E2D expressing 3xFLAG tagged R2C1 chimera (E2D:R2C1), E1DE2D expressing 3xFLAG tagged RhlE1 (E1DE2D:E1), E1DE2D expressing 3xFLAG tagged RhlE2 (E1DE2D:E2), E1DE2D expressing 3xFLAG tagged R2C1 chimera (E1DE2D:R2C1), E1E2D expressing 3xFLAG tagged R1C2 chimera (E1DE2D:R1C2).

A

```
RhlE1   1   MTFASLGLLDPLLKALEGLGHGTPTPIQAQAIPPALKGRDLLAAAQTGTGKTAGFALPLL
RhlE2   1   MSFSSLGLSEALARAVEAAGYSQPTPVQQRAIPAVLQGRDLMVAAQTGTGKTGGFALPVL

RhlE1   61  QRLTLEG-----PQVAANSVRALVLVPTRELAEQVHASVRDYGQHLPLRTAVAYGGVSIN
RhlE2   61  ERLFPAGHPDREHRHGPRQARVLVLTPTRELAAQVHDSFKVYARDLPLNSTCIFGGVGMN

RhlE1   116 PQMMKLRKGVDILVATPGRLLDLYRQNAVKFAQLQALVLDEADRMLDLGFARELDELFAA
RhlE2   121 PQIQALAKGVDVLVACPGRLLDLAGQNKVDLSHVEILVLDEADRMLDMGFIHDVKKVLAK

RhlE1   176 LPRKRQTLLFSATFSDAIRTLARELLRDPLSIEVSPRNTAAKSVRQWLVPVDKKRKAELF
RhlE2   181 LPPKRQNLLFSATFSKDIVDLANKLLHNPERIEVTPPNTTVERIEQRVFRLPAPQKRALL

RhlE1   236 CHLLQANRWRQALVFAKTRKSVEELVGLLQRQGIAADSIHGDKPQPARLRALQRFKAGEV
RhlE2   241 AHLVTVGAWEQVLVFTRTKHGANRLAEYLTKHGLPAAAIHGNKSQNARTKALADFKANDV

RhlE1   296 DLLVATDVAARGLDIEEMPLVVNFDLPIVAEDYVHRIGRTGRAGASGQAVSLVCADEVEL
RhlE2   301 RILVATDIAARGLDIDQLPHVVNYELPNVEEDYVHRIGRTGRAGRSGEAISLVAPDEEKL

RhlE1   356 LAAIETLIGQTLQRREEPDFEPEHRVPQTA---PGGVVLKKPKKPKKPKAAESVGKPGKI
RhlE2   361 LKAIEKMTRQRIPDGDAQGFDPEAVLPEVAQPEPREAPQKQPRRDKERRSSRE-RKPKDA

RhlE1   413 HLGSWFDSSAPTVKAVRKAPGFGAGAAGKPKK-RPSGK----------------------
RhlE2   420 QASNPDSNVAAAQDGTEKPAGKRRRRGGKNKENREAGQAQQPRQSREARPAKPNRPPEVD

RhlE1   449 ------------------------------------------------------------
RhlE2   480 GNRDPEEFLDDDFDNFGNRADYVSPYQGQENKGRGRRGGQQKPQGGTGQQGRGQGQGQAR

RhlE1   449 ------------------------------------------------------------
RhlE2   540 GKSQGAAQGGARGQGAGQGKAKKPRAGKPRGQGRENASRMSDAPLREPSEYGTGKQPSRQ

RhlE1   449 ----------------------------------------
RhlE2   600 PVVINKRDLVRMDRFPTAEQLDELEPRRKGERPALLTRNR
```

B

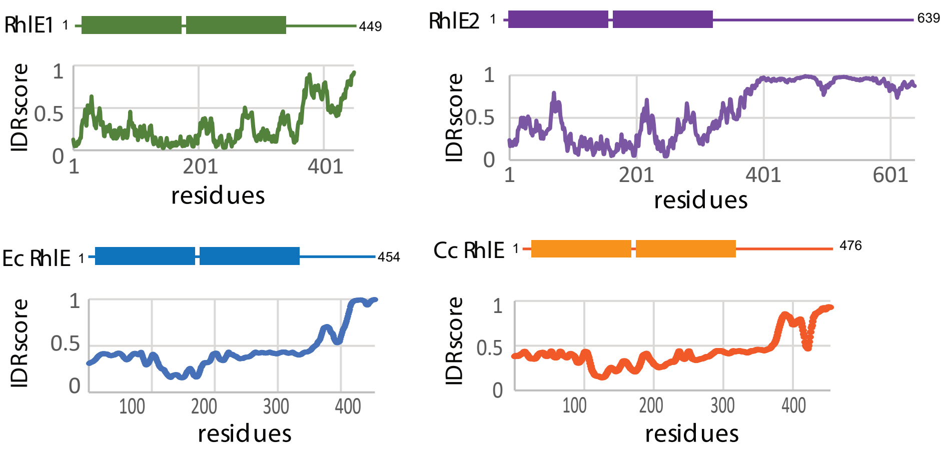


Figure S1

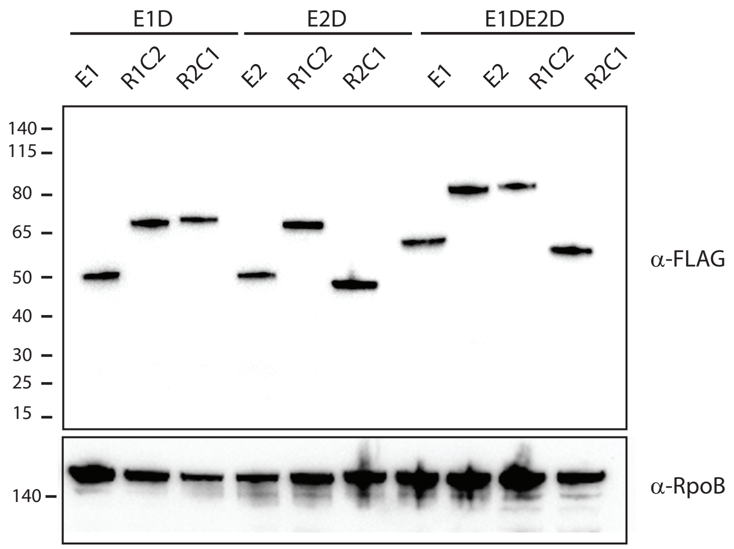
E1D
E2D
E1DE2D
E1
R1C2
R2C1
E2
R1C2
R2C1
E1
E2
R1C2
R2C1
140
115
80
65
50
40
30
25
15
α-FLAG
140
α-RpoB

Figure S2

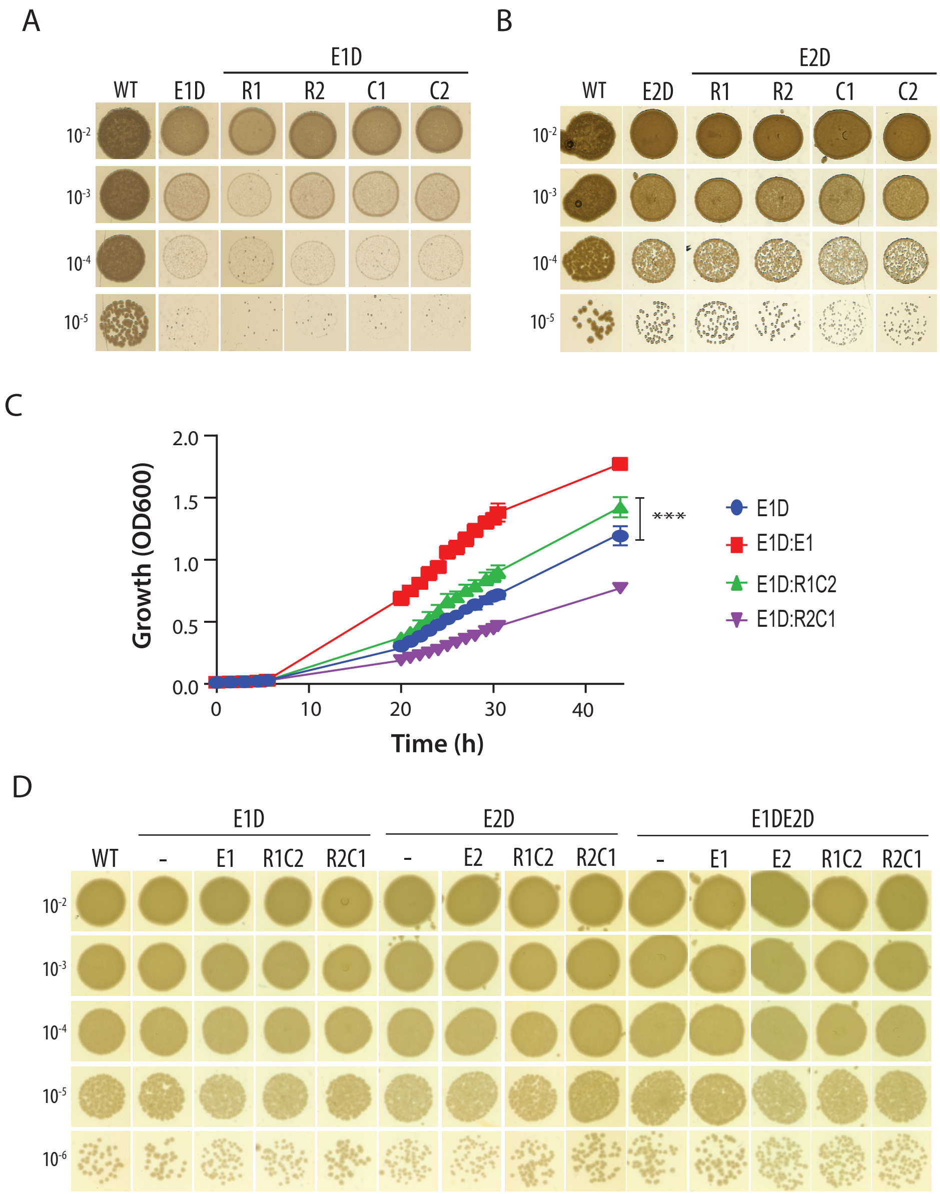
A
E1D
WT
E1D
R1
R2
C1
C2
10-2
10-3
10-4
10-5
B
E2D
WT
E2D
R1
R2
C1
C2
10-2
10-3
10-4
10-5
C
Growth (OD600)
2.0
1.5
1.0
0.5
0.0
0
10
20
30
40
Time (h)
***
E1D
E1D:E1
E1D:R1C2
E1D:R2C1
D
E1D
E2D
E1DE2D
WT
-
E1
R1C2
R2C1
-
E2
R1C2
R2C1
-
E1
E2
R1C2
R2C1
10-2
10-3
10-4
10-5
10-6

Figure S3

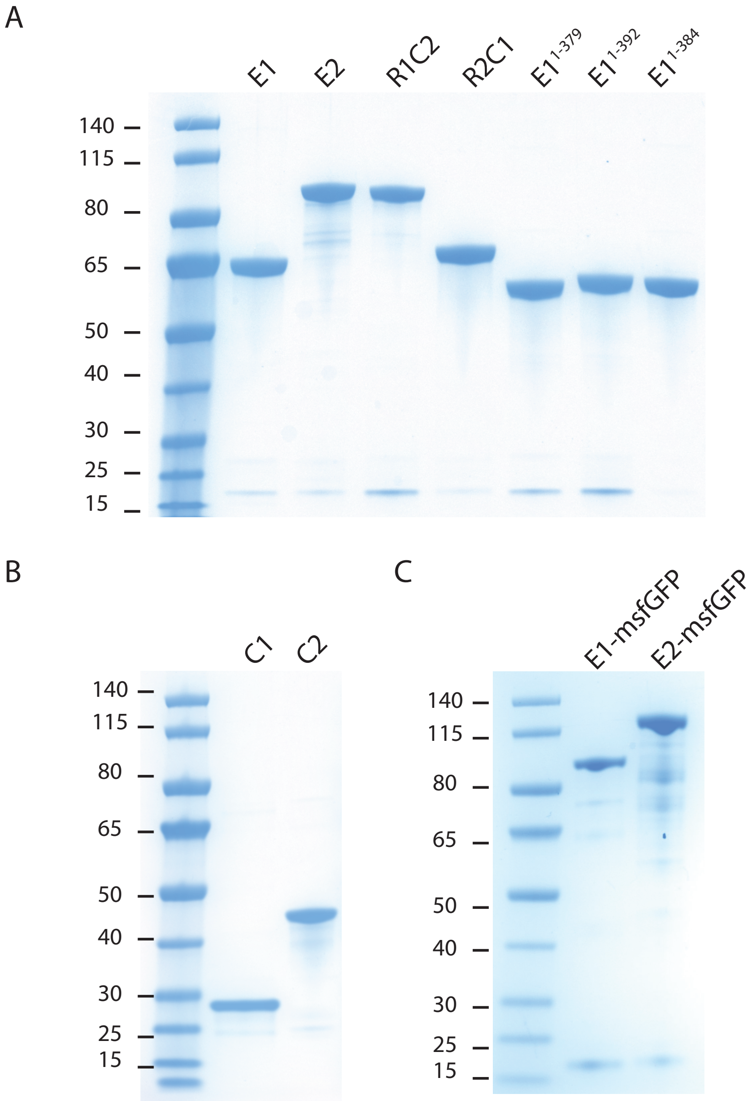
A
E1
E2
R1C2
R2C1
E1 1-379
E1 1-392
E1 1-384
140
115
80
65
50
40
30
25
15
B
C1
C2
140
115
80
65
50
40
30
25
15
C
E1-msfGFP
E2-msfGFP
140
115
80
65
50
40
30
25
15

Figure S4

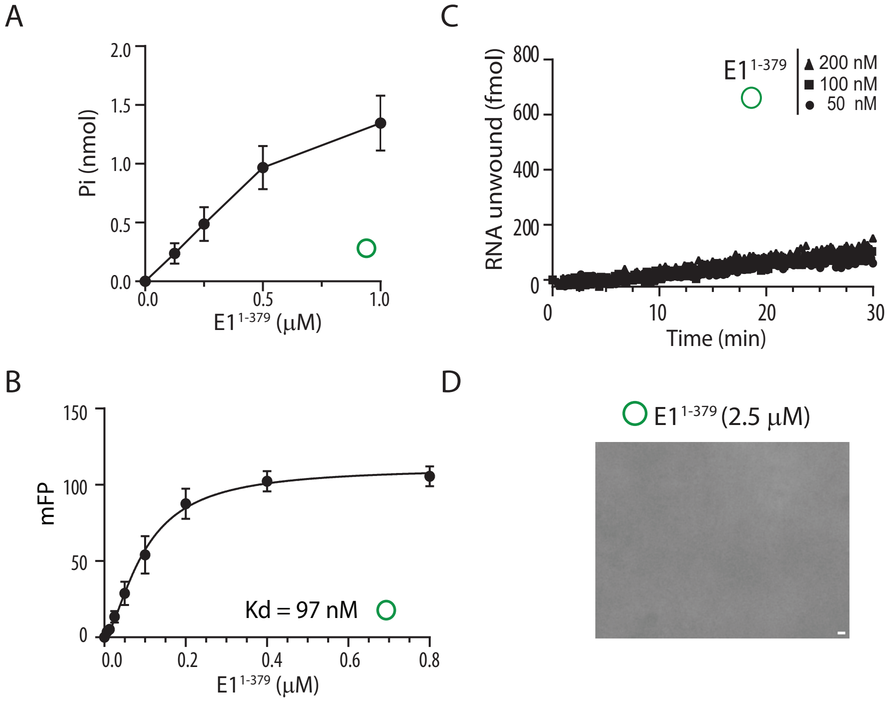
A
Pi (nmol)
2.0
1.5
1.0
0.5
0.0
0.0
0.5
1.0
E1 1-379 (μM)
B
mFP
150
100
50
0
0.0
0.2
0.4
0.6
0.8
Kd = 97 nM
E1 1-379 (μM)
C
RNA unwound (fmol)
800
600
400
200
0
E1 1-379
200 nM
100 nM
50 nM
0
10
20
30
Time (min)
D
E1 1-379 (2.5 μM)

Figure S5

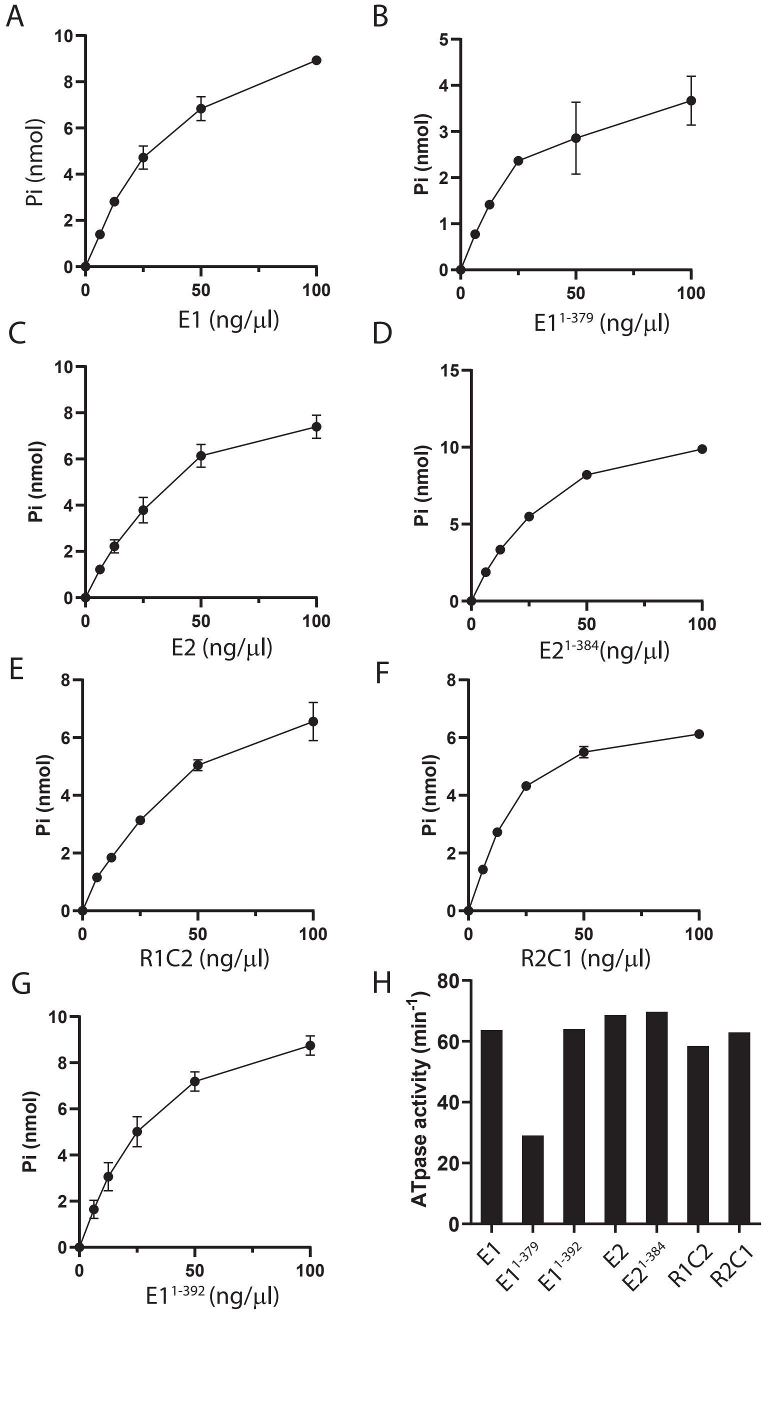
A
Pi (nmol)
E1 (ng/μl)
B
Pi (nmol)
E1 1-379 (ng/μl)
C
Pi (nmol)
E2 (ng/μl)
D
Pi (nmol)
E2 1-384(ng/μl)
E
Pi (nmol)
R1C2 (ng/μl)
F
Pi (nmol)
R2C1 (ng/μl)
G
Pi (nmol)
E1 1-392 (ng/μl)
H
ATpase activity (min-1)
E1
E1 1-379
E1 1-392
E2
E2 1-384
R1C2
R2C1
0
50
100

Figure S6

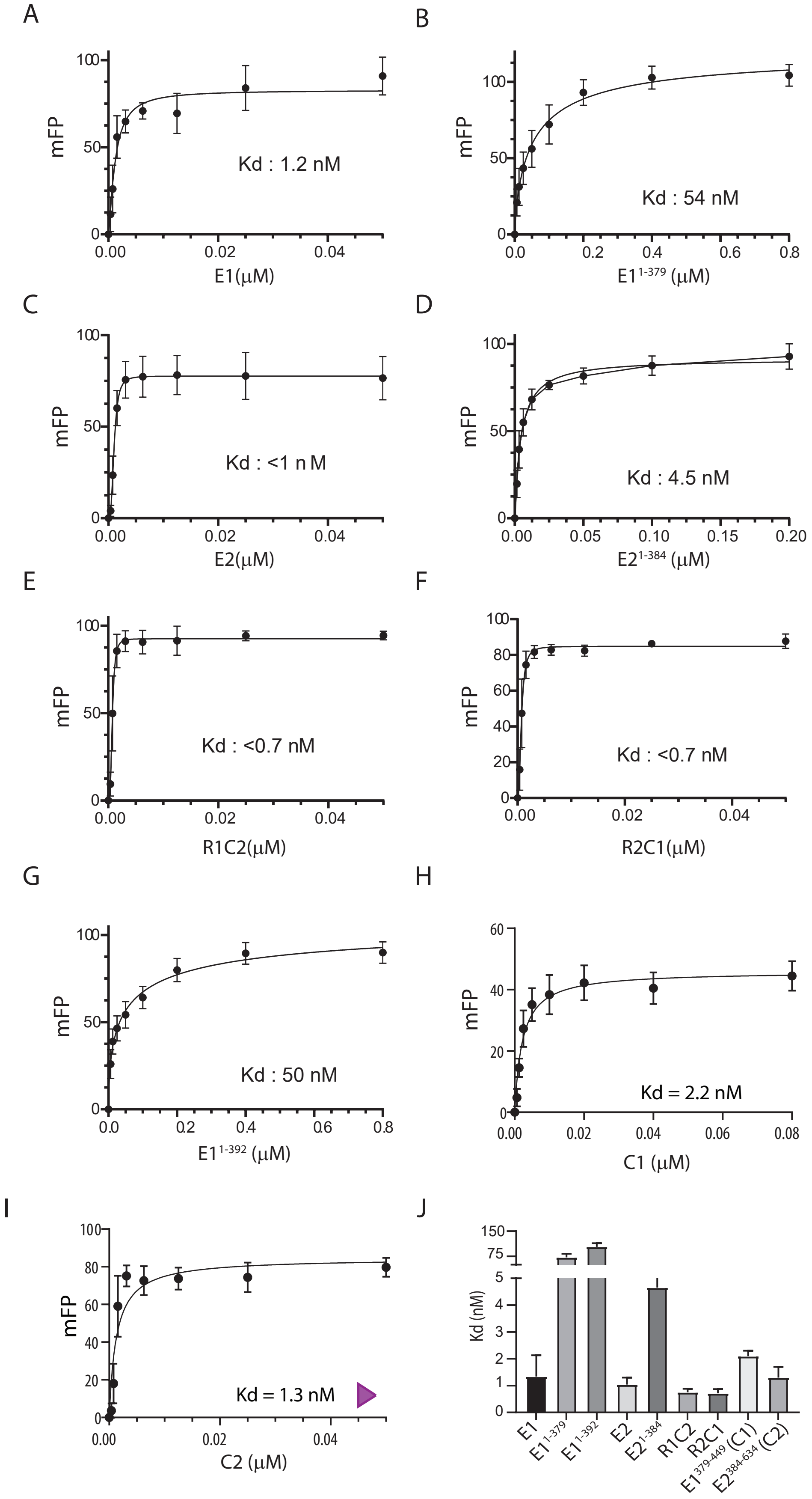
A
mFP
Kd : 1.2 nM
E1(μM)
B
mFP
Kd : 54 nM
E1 1-379 (μM)
C
mFP
Kd : <1 n M
E2(μM)
D
mFP
Kd : 4.5 nM
E2 1-384 (μM)
E
mFP
Kd : <0.7 nM
R1C2(μM)
F
mFP
Kd : <0.7 nM
R2C1(μM)
G
mFP
Kd : 50 nM
E1 1-392 (μM)
H
mFP
Kd = 2.2 nM
C1 (μM)
I
mFP
Kd = 1.3 nM
C2 (μM)
J
Kd (nM)
E1
E1 1-379
E1 1-392
E2
E2 1-384
R1C2
R2C1
E1 379-449 (C1)
E2 384-634 (C2)

Figure S7

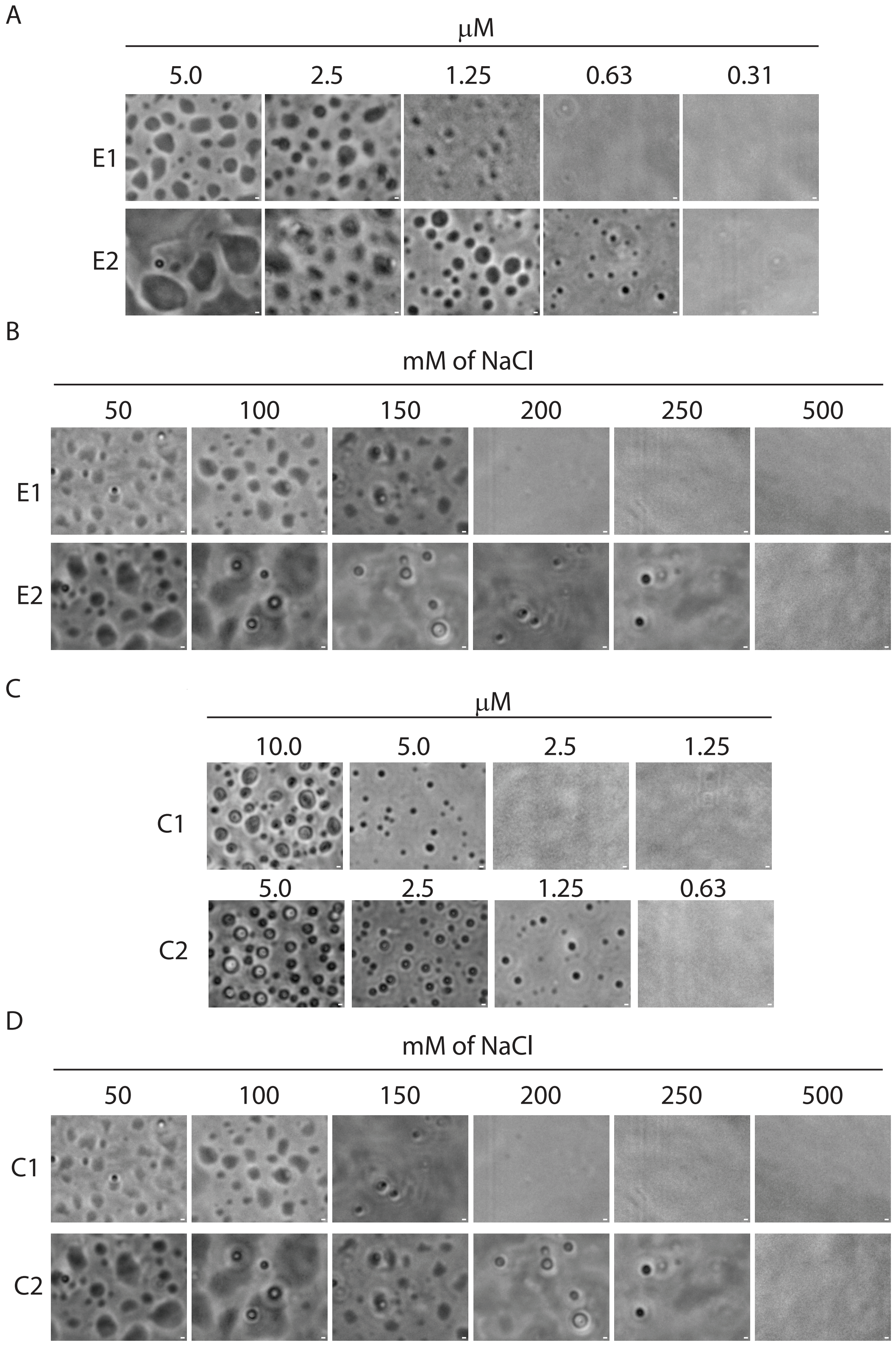
A
μM
5.0
2.5
1.25
0.63
0.31
E1
E2
B
mM of NaCl
50
100
150
200
250
500
E1
E2
C
μM
10.0
5.0
2.5
1.25
C1
5.0
2.5
1.25
0.63
C2
D
mM of NaCl
50
100
150
200
250
500
C1
C2

Figure S8

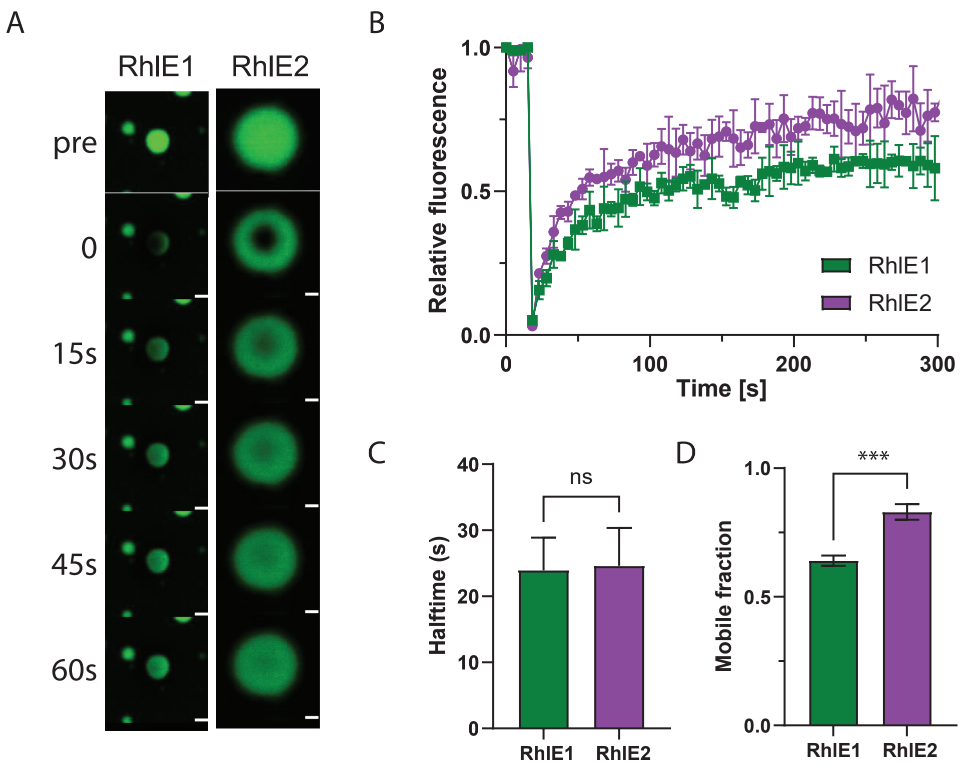
A
RhlE1
RhlE2
pre
0
15s
30s
45s
60s
B
Relative fluorescence
1.0
0.5
0.0
0
100
200
300
Time [s]
RhlE1
RhlE2
C
Halftime (s)
40
30
20
10
0
ns
RhlE1
RhlE2
D
Mobile fraction
1.0
0.5
0.0
***
RhlE1
RhlE2

Figure S9

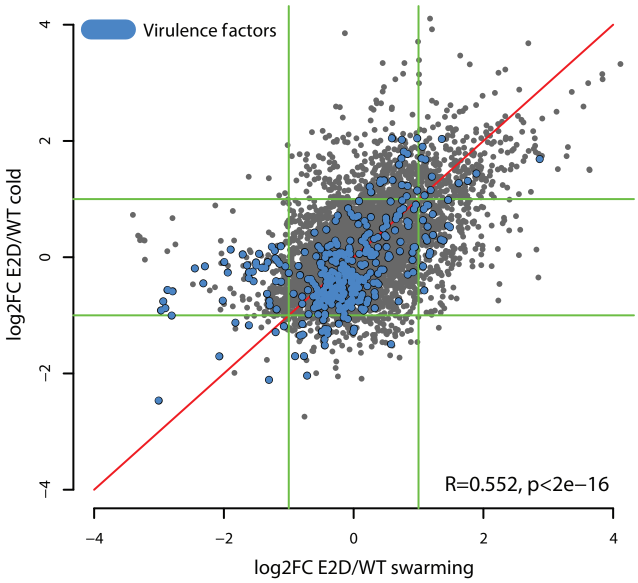

Virulence factors
log2FC E2D/WT cold
log2FC E2D/WT swarming
R=0.552, p<2e−16
−4
−2
0
2
4


Figure S10

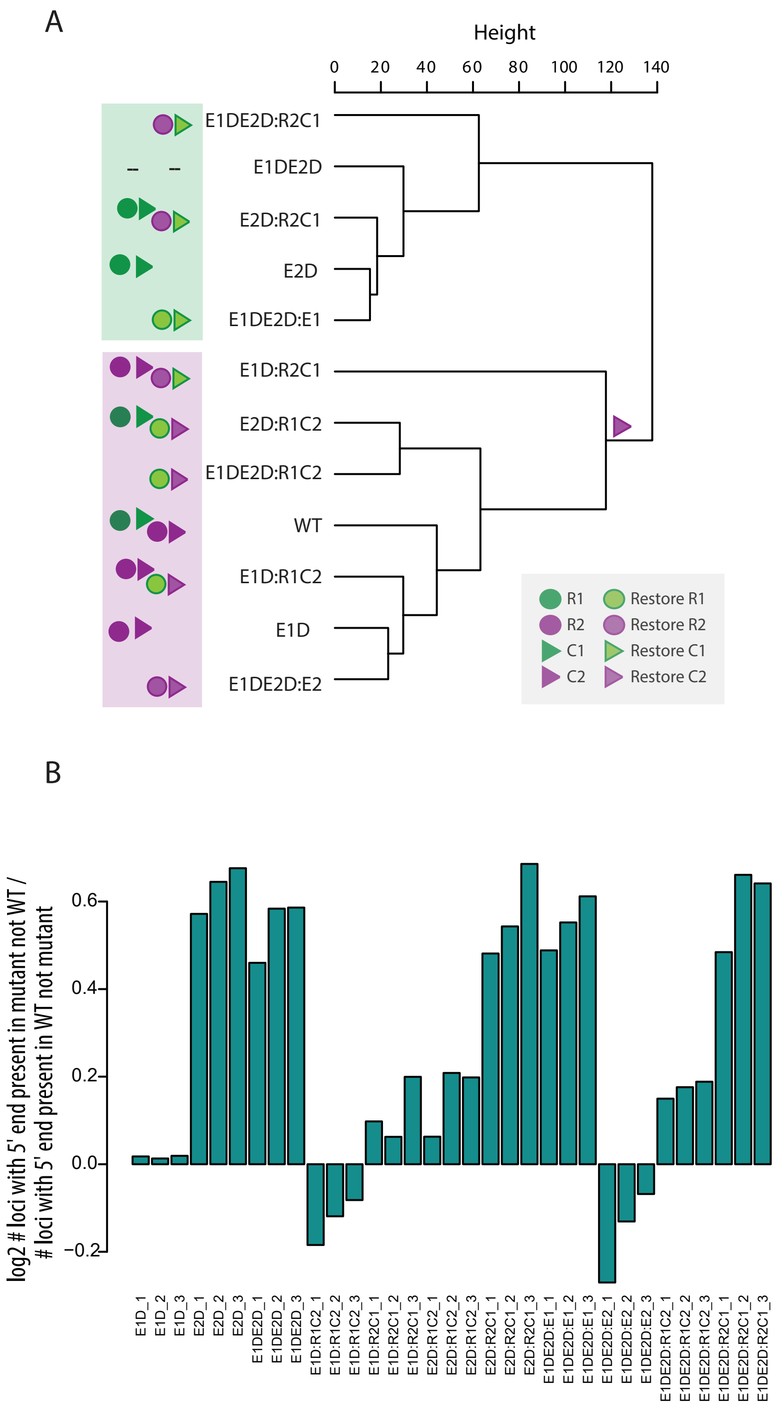
A
Height
0 20 40 60 80 100 120 140
E1DE2D:R2C1
E1DE2D
E2D:R2C1
E2D
E1DE2D:E1
E1D:R2C1
E2D:R1C2
E1DE2D:R1C2
WT
E1D:R1C2
E1D
E1DE2D:E2
R1
Restore R1
R2
Restore R2
C1
Restore C1
C2
Restore C2
B
log2 # loci with 5' end present in mutant not WT / # loci with 5' end present in WT not mutant
0.6
0.4
0.2
0.0
−0.2
E1D_1
E1D_2
E1D_3
E2D_1
E2D_2
E2D_3
E1DE2D_1
E1DE2D_2
E1DE2D_3
E1D:R1C2_1
E1D:R1C2_2
E1D:R1C2_3
E1D:R2C1_1
E1D:R2C1_2
E1D:R2C1_3
E2D:R1C2_1
E2D:R1C2_2
E2D:R1C2_3
E2D:R2C1_1
E2D:R2C1_2
E2D:R2C1_3
E1DE2D:E1_1
E1DE2D:E1_2
E1DE2D:E1_3
E1DE2D:E2_1
E1DE2D:E2_2
E1DE2D:E2_3
E1DE2D:R1C2_1
E1DE2D:R1C2_2
E1DE2D:R1C2_3
E1DE2D:R2C1_1
E1DE2D:R2C1_2
E1DE2D:R2C1_3

Figure S11
